# Computational Design of a Highly-Specific HVEM-Based Inhibitor of LIGHT Protein

**DOI:** 10.1101/2023.10.10.561657

**Authors:** Piotr Ciura, Pamela Smardz, Magdalena A. Krupa, Marta Spodzieja, Pawel Krupa, Adam K. Sieradzan

## Abstract

**Motivation:** HVEM-LIGHT binding regulates the immune system response in various ways: it co-stimulates T cell proliferation; promotes B cell differentiation and secretion of immunoglobulins; and enhances dendritic cell maturation. Strong and prolonged stimulation of T cells to proliferate causes high levels of IFN–*γ*, which leads to chronic inflammation and is the reason for various autoimmune diseases. Therefore, blocking HVEM-LIGHT interaction may be a way to cure these diseases and prevent adverse reaction in organ and tissue transplantation.

**Results:** In this work, we designed 62 peptides based on the CRDs of the HVEM structure, differentiating in the number and combination of disulfide bonds present. Based on extensive all-atom MD simulations in state-of-the-art force fields, followed by MM-GBSA binding energy estimation, we selected the most promising CRD2 variants interacting with LIGHT. Several point mutations of these variants provided us with the most strongly binding moiety: the CRD2 with a single disulfide bond (C58-C73) and K54E substitution. This result was supprased only by the truncated variants of CRD2(39-73) with the same disulfide bond present. The binding mechanism was investigated by the use of steered MD simulations, which showed the increased binding affinity of the abovementioned variants, while experimental circular dichroism was used to determine their structural properties.

**Availability and Implementation:** Three PDB models of the LIGHT inhibitors: PM0084527, PM0084528, and PM0084592.

**Supplementary information:** Online supplementary data is available at: .

## Introduction

The herpesvirus entry mediator (HVEM) and its ligand LIGHT are members of the tumor necrosis factor superfamily (TNFSF) (Harrop *et al*., 1998a; Mauri *et al*., 1998). These proteins play multiple roles in the immune system by regulating diverse processes and T cell responses. In particular, HVEM and LIGHT function as co-stimulatory molecules, and their interaction results in the enhancement of T-cell growth, differentiation, and cytokine secretion (Wang *et al*., 2001; Tamada *et al*., 2000). HVEM is a type I transmembrane glycoprotein composed of 283 amino-acid residues, divided into a signaling peptide (-37-0) and extracellular (1-164), transmembrane (165-185), and cytoplasmic (186-245) parts. The extracellular fragment of HVEM contains four cysteine-rich domains (CRD1: 1-38, CRD2: 39-81, CRD3: 82-124, CRD4:125-164), and the disulfide bonds within these domains are critical for establishing the tertiary structure of this receptor (Marsters *et al*., 1997; Kwon *et al*., 1997). LIGHT, also known as TNFSF14, is a homotrimeric type II transmembrane protein expressed on activated T cells and immature dendritic cells (Mauri *et al*., 1998; Tamada *et al*., 2000). It is involved in various biological processes, including inflammation, immune regulation, and apoptosis, and plays a critical role in the pathogenesis of various autoimmune diseases and cancer (Ware, 2009), as well as, COVID-19 induced pneumonia and inflammations (Ware *et al*., 2022). Therefore, understanding the molecular mechanisms of LIGHT signaling is crucial for developing new therapies for these diseases. HVEM provides a stimulatory signal by interacting with the LIGHT and LT*α* ligands via the CRD2 and CRD3 domains (Mauri *et al*., 1998; Marsters *et al*., 1997) (Fig. 1), and an inhibitory signal when it binds to the BTLA (Sedy *et al*., 2005) or CD160 (Cai *et al*., 2008) receptors via the CRD1 domain. On the other hand, neither the CRD4 structure nor its affinity to other molecules is known. (Liu *et al*., 2021). This binding pattern of HVEM offers multiple targetable molecules for modulating immunological responses. Moreover, the HVEM-LIGHT complex could be involved in various inflammatory processes such as inflammatory bowel disease (Cohavy *et al*., 2005) and rheumatoid arthritis (RA) (Kim *et al*., 2005; Pierer *et al*., 2007) due to the modulation of T cell proliferation. Furthermore, studies suggest that the interaction between these proteins may play a major role in transplantation. For example, targeting LIGHT protein using HVEM-Ig and LT*β*R-Ig fusion proteins significantly reduces allogenic T cell immune responses, including proliferation and cytotoxic T cell activity (Tamada *et al*., 2000). Similarly, the use of HVEM mAbs has shown a similar effect (Harrop *et al*., 1998b). All these data highlight that the HVEM-LIGHT complex could be a key factor in controlling T cell immune responses and may be a candidate to target in different immune-mediated diseases.

**Fig. 1:**
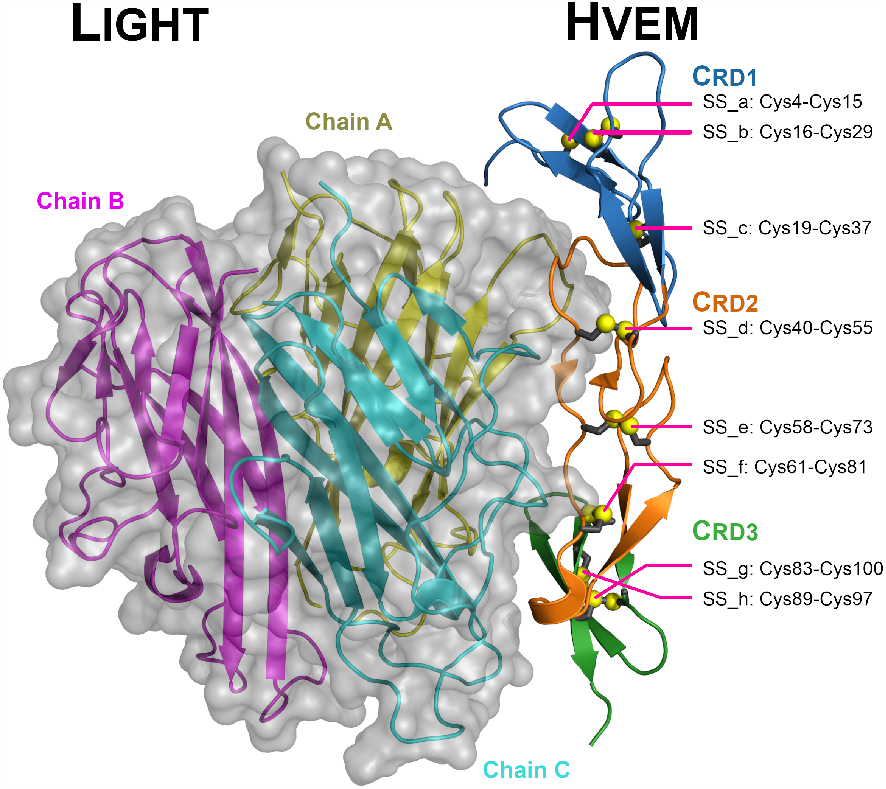
Cartoon representation of the LIGHT-HVEM complex with the semitransparent surface representation of the LIGHT trimer (PDB code: 4RSU). CRD1, CRD2 and CRD3 are indicated as blue, orange, and green colors, respectively, while, chains A, B and C from LIGHT trimer are colored yellow, magenta and cyan respectively. HVEM disulfide bonds are shown as grey sticks and yellow spheres (a-h).

As short peptides usually have lower toxicity, drug-drug interaction and production cost (Diao and Meibohm, 2013), the aim of the present study is to propose short peptide inhibitors of the HVEM-LIGHT interaction based on HVEM-binding fragments. To achieve this, we designed peptides that are fragments of HVEM molecule, which can potentially block the formation of the HVEM-LIGHT complex and determined their affinity to the trimeric LIGHT protein. We performed detailed analyses of binding free energies, dynamics, and stability of the obtained complexes based on extensive molecular dynamics (MD) simulations. This method proved to be an effective tool for peptide design in other HVEM ligand systems, those peptides proved to bind strongly in vitro (Kuncewicz *et al*., 2023). We also determined amino-acid residues and disulfide bonds that have a crucial influence on the formation of the HVEM-LIGHT complex and identified the binding mechanism to LIGHT for the designed peptides. This allowed us to obtain a single-residue mutant of the CRD2 domain (CRD2e_K54E) with a single disulfide bond (C58-C73) that binds most strongly to LIGHT out of all the examined variants based on whole domains and the truncated variants of the CRD2 domain (CRD2(39-73)e) with the same disulfide bond present, which are characterized by even stronger binding, despite being shorter in length. These results were further confirmed by the steered MD (SMD) simulations.

## Methods

### Simulated systems

Our studies involved MD simulations of 62 molecules, based on the HVEM protein, including various lengths, disulfide-bond combinations and amino-acid substitutions interacting with the LIGHT trimer (Table S1), while selected 24 variants were simulated also alone, in bulk water, without the presence of LIGHT protein, for sake of the comparison. To simplify the naming, each disulfide bond was assigned a letter from a to h (Fig. 1), based on the position of the first cysteine residue involved in a bond (detailed explanation is provided in Table S1).

### All-atom Molecular Dynamics simulations

The first set of MD simulations was used to determine the stability of the HVEM protein and its variants in an aquatic environment, while the second set was used to determine the binding affinity of proposed inhibitors to LIGHT and the change of structural properties upon binding. All MD simulations adapted the experimental model of HVEM-LIGHT (PDB: 4RSU, (Liu *et al*., 2021)) as the initial conformation, with or without disulfide-bond modifications, amino-acid substitutions and truncations (Table S1).

All systems were prepared using the tleap program with the ff19SB force field (Tian *et al*., 2020) and four-point OPC water model (Izadi *et al*., 2014), while the simulations were run in pmemd.cuda, part of Amber22 (Case *et al*., 2022). The simulation details are described in SI section **All-atom Molecular Dynamics simulations details**.

In addition to classical MD simulations, of which 11 lowest energy variants determined by MM-GBSA we gathered in Fig. 2, a series of SMD simulations (25 trajectories per system, each of about 80ns) was run to establish a binding-unbinding mechanism for CRD2e, CRD2e_K54E, and CRD2(39-73)e (Fig. 3) representative models.

**Fig. 2:**
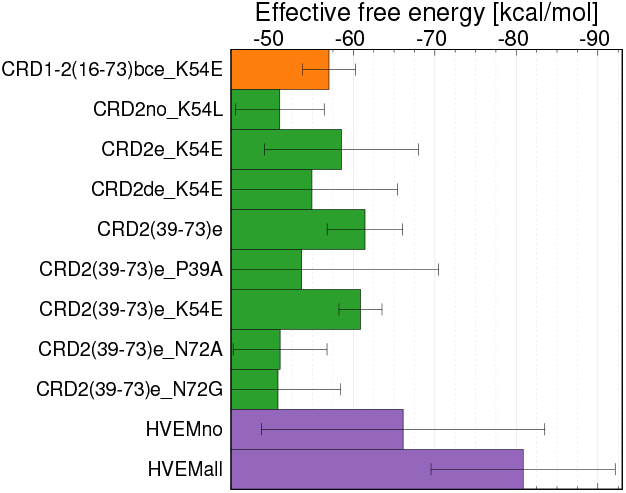
Bar plot of the effective binding free energy for HVEM variants with LIGHT with the highest binding affinity (*<* −50kcal/mol).

**Fig. 3:**
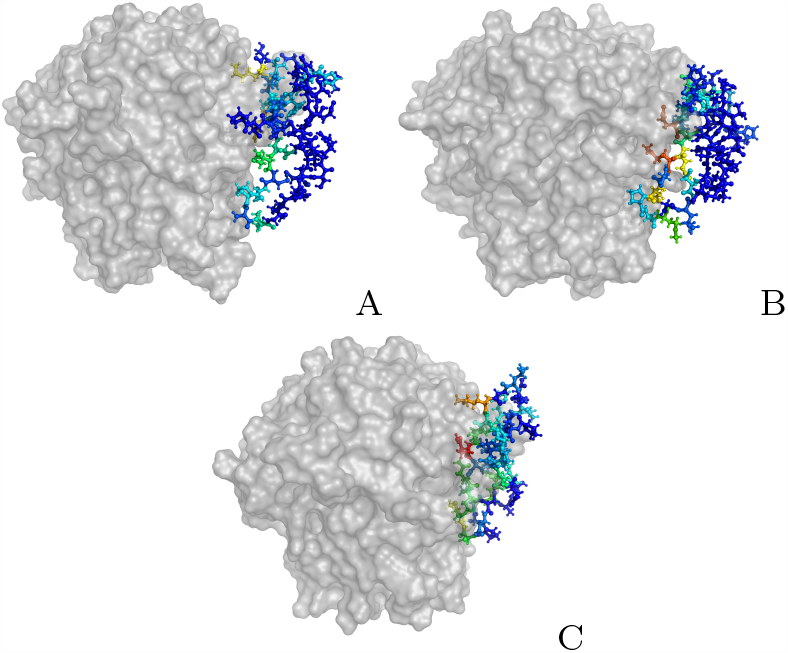
Ball-and-stick representation of HVEM-based variants colored by the pairwise effective binding energy (rainbow colors from blue to red, where blue indicates the lowest binding energy and red is the highest) for: A) CRD2e, B) CRD2e_K54E, and C) CRD2(39-73)e interacting with the LIGHT trimer (grey surface).

### Analysis

All analyses were performed on the converged parts of the trajectories: the last 20 (out of 200 ns) and 500 (out of 1000 ns) for simulations of HVEM with and without LIGHT, respectively, and averaged out over three independent trajectories. The free energy change between the bounded and free states of the receptor-ligand complexes was estimated using the Molecular Mechanics - Generalized Born Surface Area (MM-GBSA) method (Wang *et al*., 2019) with the recommended radii set and GB estimation method (mbondi3 and igb=8) (Nguyen *et al*., 2013). Such combination of force-field, water model and MM-GBSA method is the modern standard for the binding affinity prediction (Chen and Zacharias, 2023). Since calculating entropic effects is computationally expensive and may significantly vary between selected snapshots, the effective free energy (Liu and Beveridge, 2002) was analyzed, which should provide good qualitative results for similar compounds (Fatima *et al*., 2023). Energy decomposition was performed on per-residue and pairwise per-residue basis, with the latter summarized for each residue in the HVEM variant of its interactions only with LIGHT residues, to show the difference in how each residue influences overall binding stability and the influence on the interaction partner.

Analyses of: C*α* root-mean-square deviation (RMSD); root-mean-square fluctuations (RMSF); radius of gyration (RG); maximum distance of any heavy atom to center of mass of the protein (RGmax); solvent-accessible surface area (SASA) (using the LCPO algorithm (Weiser *et al*., 1999)); contacts based on distance cutoffs of 8 °A and 6°A for determining dissociation of the complex in MD and SMD, respectively, (contact is defined as 0 if there are no heavy atoms from the complex partner within the range, and as 1 if there is at least one such heavy atom); fraction of secondary structure with DSSP algorithm (Kabsch and Sander, 1983) implemented in AmberTools2022 cpptraj(Case *et al*., 2022). For comparison, the fraction of secondary structure was analyzed with STRIDE (Heinig and Frishman, 2004) and KAKSI (Martin *et al*., 2005). LRMSD, calculated only for the backbone of the ligand after superposition on the receptor, was determined using DockQ (Basu and Wallner, 2016). As STRIDE, KAKSI and DockQ can process only a single structure at the time, an in-house script was used to process equilibrated parts of trajectories. Representative models were generated by hierarchical clustering of the converged parts of the simulation. All figures were prepared with the use of Python 3.7 and gnuplot5.2, while structures were visualised with PyMOL 2.5.0 (Schrödinger and DeLano, 2021).

### Peptide synthesis, purification, and circular dichroism spectra measurement

Selected peptides were synthesized using the solid-phase peptide synthesis (SPPS) method and subsequently purified using reverse-phase high-performance liquid chromatography (RP-HPLC) and then disulfide bonds were formed using a designed procedure - a detailed description of these steps is available in the SI section **Peptide synthesis; Peptide purification; Disulfide bond formation**.

Circular dichroism (CD) spectra were recorded at the Circular Dichroism Laboratory, Faculty of Chemistry, University of Gdańsk. The analysis was carried out using a CD J-815 circular dichroism spectrometer by Jasco. Peptide solutions with concentrations of 0.15 mg/mL were used for the tests, and all CD spectra were taken in water at 298K. The results are presented in the form of the dependence of the molar ellipticity on the wavelength.

### Selectivity check

To check if the designed peptides characterized by the strongest effective binding energy are binding selectively to the LIGHT binding pocket, docking studies with use of HDOCK(Yan *et al*., 2020) were performed. We also checked if the molecules can effectively bind to other protein, which in physiological conditions binds the HVEM protein, BTLA.

## Results

### Structural properties of unbound HVEM protein and its disulfide-bond variants

For all of the structural fragments in which cysteine residues forming disulfide bonds are present, reduction of it causes spatial separation of these elements (Fig. S1). Only in the whole HVEM simulations, CRD2 maintain close native contacts between reduced cysteins. In general, the largest and lowest structural fluctuations were observed in all HVEM domains when all disulfide bonds were absent and present, respectively (Fig. S1). The introduction of any single disulfide bond consistently enhances structural stability. However, the strengthening of this effect does not always occur with the addition of another disulfide bond. Further structural stabilization is observed when the rest of the HVEM molecule is present. This phenomenon is particularly pronounced in the case of CRD2, which exhibits remarkable stability even in the absence of any disulfide bonds when surrounded by CRD1 and CRD3.

In CRD2, which is the most important domain for binding LIGHT, the disulfide bond between Cys58-Cys73 (e) is the only one that significantly increases structural stability on its own (reducing RMSD from 10.52*±*0.28 to 4.93 *±* 0.65°A) and can be compared to the stability of the two-disulfide bonds variants CRD2df and CRD2ef. Only when combined with Cys40-Cys55 its stability slightly increases (CRDde: 3.84 *±* 0.79 °A), approaching the case when all three bonds are present (CRD2def: 2.60 *±* 0.18° A).

The radius of gyration (RG) and RGmax analyses shows that the compactness of CRD2 is not significantly affected by the presence or absence of disulfide bonds, except for CRD2no and CRD2d (Fig. S2), which exhibited a more relaxed or loose structure.

The details of influence of all disulfide bonds on CRD stability is described in the SI, section **Influence of disulfide bonds on CRD stability** (Fig. S1).

Content of both *α*-helices and *β*-sheets in HVEM domains is heavily impacted by the presence or absence of given disulfide bonds and the rest of the HVEM polypeptide chain (Table S2). While full HVEM sequence has a strong tendency to form *β*-sheets over *α*-helices, which is understandable, as this secondary structure element requires distant parts of the protein to obtain complete stabilization, domains usually prefer more disordered and *α*-helical structures. The largest differences are observed for variants with all and no of the disulfide-bonds present, however, a truncation of the CRD2 further modify its behavior. Despite that, the predominant type of the secondary structure in all of the HVEM variants is always disordered (coil) and turn, which is closely followed by *β*-sheets, while *α*-helices content is always low (*<* 8%). Addition of the disulfide bonds have various degree of impact, but predominantly draws the secondary-structure content close to the all-disulfide-bond variants and the complete HVEM chain with all disulfide bonds present.

The CD spectra generated with web-server (Mavridis and Janes, 2016) reveal some helical (Fig. S3B) content which is not present in analysis with other methods and is not observed in the experimental CD spectra (Fig. S3A). Despite the fact that CD is a very low-resolution method of secondary structure determination, especially for flexible structures,(Nagy *et al*., 2019) for all peptides coil structure is dominant one as the value of molar ellipticity is below 0 for 200 nm, which in agreement with all secondary structure determination tools used for this simulations. Therefore, it can be concluded that peptides do not form significant amounts of stable secondary structures, and are mainly unstructured.

### Changes of the structural properties of HVEM and its variants upon binding LIGHT trimer

HVEM variant structures become more rigid upon LIGHT binding, which is particularly evident in variants without any disulfide bonds present. This effect is even more pronounced when the complex exhibits high binding affinity (Fig. S2). However, upon binding, the structure of CRD2 variants becomes slightly more expanded. This is evident in the relatively small increase in the RG and a larger increase in RGmax, with the only exception being CRD2no. In the case of CRD2no, the combination of high flexibility and high binding affinity makes it more strongly impacted by the binding of LIGHT. Interestingly, the theoretical SASA of HVEM variants, calculated without the presence of any other molecules, remains similar for both free and bound HVEM variants with only slight increase (Fig. S4). This observation suggests that hydrophobic interactions are not the primary contributors to HVEM-LIGHT interactions.

In general, binding of HVEMall to LIGHT does not have a significant impact on the fluctuations (RMSF; Fig. S5) and secondary structure of the molecule (Fig. S6). Moreover, it’s noteworthy that the most structurally stable region upon binding to LIGHT is CRD2. The most stable HVEM variant overall is CRD2all, which secondary structure remains unchanged if all disulfide bonds are present, similarly to the complete HVEM chain with all disulfide bonds present. The most drastic changes of the secondary structure upon LIGHT binding are observed for CRD3no, which shifts strongly into the *β*-sheets.

### Effective binding energy to select the best candidates for LIGHT inhibition

The effective binding energy of HVEM and its variants to the LIGHT trimer was determined using the MM-GBSA method, while the contribution of particular amino-acid residues to the total effective binding energy was calculated using per-residue and pairwise energy decomposition, however, only for the complexes which did not dissociate during MD simulations (Fig. S7). Our analysis revealed that, among the three HVEM domains, CRD2 emerged as the primary and the robust interacting entity with the LIGHT trimer (Fig. S8-9), therefore, our further efforts to design the strongest-binding peptide was focused mostly on this HVEM fragment. It should be noted that disulfide bonds have tremendous influence on the binding affinities, i.e. CRD2no (CRD2 domain without disulfide bonds) demonstrated significantly higher affinity to LIGHT compared to CRD2all (with all disulfide bonds).

In the next step, we established that Lys54 has the most unfavorable energetic effect in most of the CRD2 variants; therefore, it was our first candidate for point mutations in order to attempt to increase binding affinity to the LIGHT trimer. Serine, leucine, isoleucine, glutamic acid, aspartic acid, tyrosine, and valine were selected as candidates for replacements to scan amino-acid residues of the opposite charge and properties to lysine. We selected CRD2 variants with different disulfide bonds pattern which exhibited the best affinity: CRD2no and CRD2de, and the worst, but most stable in solution, one: CRD2e.

Unfortunately, no point mutations in CRD2no significantly improved the binding energy, while the mutation to serine dramatically reduced the binding affinity (-21.46 *±* 3.90 kcal/mol). In the case of CRD2de, only the mutation to glutamic acid improved the effective binding energy (-54.92 *±* 10.52 kcal/mol). Surprisingly, the highest effective binding energy was observed for the same substitution but in the CRD2e variant, which is characterized by the lowest binding affinity among all CRD2 variants but with the single-point mutation becomes the one with the highest affinity (CRD2e_K54E: -58.55 *±* 9.45 kcal/mol). Smaller improvement was observed for CRD2e with a substitution of Lys54 to valine (-41.17 *±* 7.20 kcal/mol).

We also designed a series of double-point mutants to mitigate the unfavorable energy effect from Asp62, which was emphasized in the designed mutants. This residue was replaced with alanine, leucine, lysine, and serine. Unfortunately, none of the designed double mutants exhibited a more favorable effective binding energy than the original variant (Fig. S8).

Based on the binding energy decomposition (Fig. S9), we determined that not all amino-acid residues are involved in the formation of a complex with the LIGHT trimer. Therefore, we designed four peptides accordingly: two based on the CRD1 domain, namely CRD1(16-38)bc and CRD1(16-38)no, and two on the CRD2 domain - CRD2(39-73)e and CRD2(39-73)no, which included only amino-acid residues with sufficient affinity for LIGHT trimer. It should be noted that, in general, peptides based on the CRD1 domain have much lower affinity for LIGHT than those based on the CRD2 domain, and the truncated variant showed even lower affinity (-18.89 *±* 6.41 and -44.35 *±* 6.06 kcal/mol respectively). However, truncation of the CRD2 amino-acid residues resulted in a significant increase in effective binding energy, with CRD2(39-73)e peptide being the strongest binding peptide (-61.43 *±* 4.63 kcal/mol), despite its shorter polypeptide chain. Conversely, CRD2(39-73)no exhibited lower affinity than CRD2no. Remarkably, the removal of the 8 C-terminal residues from CRD2e resulted in a CRD2(39-73)e conformation that barely extends beyond the binding site of the LIGHT trimer (Fig. 3).

**Table 1.** Top 5 amino-acid residues, from CRD2e, CRD2e_K54E, and CRD2(39-73)e and LIGHT protein in complexes, based on their effective binding free energy decomposition [kcal/mol] determined by MM-GBSA pairwise analysis.

| CRD2e | CRD2e_K54E | CRD2(39-73)e |
| --- | --- | --- |
| Gln57 $-9.04 \pm 3.67$ | Met60 $-11.76 \pm 2.94$ | Gln57 $-13.90 \pm 2.70$ |
| Lys54 $-8.09 \pm 4.20$ | Gln57 $-11.34 \pm 3.99$ | Lys54 $-10.21 \pm 8.46$ |
| Met60 $-4.74 \pm 2.15$ | Asp62 $-8.48 \pm 3.60$ | Met65 $-8.01 \pm 3.62$ |
| Met65 $-3.87 \pm 3.11$ | Gln59 $-7.94 \pm 2.34$ | Arg68 $-6.03 \pm 5.10$ |
| Gln59 $-2.95 \pm 1.86$ | Met65 $-6.28 \pm 2.03$ | Gln59 $-5.87 \pm 2.86$ |
| LIGHT | LIGHT | LIGHT |
| Asn160 $-5.37 \pm 2.26$ | Arg286 $-9.61 \pm 2.15$ | Glu87 $-9.84 \pm 3.43$ |
| Arg286 $-5.08 \pm 2.27$ | Asn160 $-8.18 \pm 1.91$ | Gly177 $-5.08 \pm 1.53$ |
| Tyr82 $-5.03 \pm 1.21$ | Arg284 $-5.26 \pm 2.19$ | Arg81 $-5.01 \pm 2.85$ |
| Asp287 $-3.42 \pm 2.26$ | Arg81 $-4.71 \pm 2.50$ | Arg286 $-4.84 \pm 1.01$ |
| Glu84 $-3.14 \pm 1.64$ | Ser161 $-4.18 \pm 1.60$ | Asp287 $-4.34 \pm 1.71$ |

In an effort to further improve the affinity of CRD2(39-73)e, we designed a series of mutants in which the Pro39, Lys54, Cys61, and Asn72 residues were substituted by other amino-acid residues. Unfortunately, these mutants displayed lower affinity for the LIGHT trimer than the parent peptide.

Our final attempt was to design a peptide combining the best CRD1 and CRD2 variants, namely CRD1(16-38)bc and CRD2(39-73)e peptides. Its affinity for the LIGHT protein, although higher than peptide comprising only the fragment of the second domain (-49.53 *±* 3.04 kcal/mol compared to CRD2e -35.26 *±* 1.94 kcal/mol), doesn’t show significant improvement compared to CRD2(39-73)e and CRD2e_K54E. Therefore we decided to try the K54E mutation, which was determined to improve affinity in the case of CRD2 peptides. A mutant designed this way, CRD1-2(16-73)bce K54E, exhibited an effective free energy slightly lower than the CRD2e_K54E peptide. Due to our desire to design the shortest possible peptides, further research on this compound was abandoned.

### Binding energy and process captured by the SMD simulations

To obtain a more complete view of the interactions influencing the binding between HVEM variants and LIGHT trimer, a series of 25 SMD trajectories for representative conformations of the three most promising systems were performed, namely CRD2, CRD2e_K54E, and CRD2(39-73)e, in which the complex partners were extended from each other by using a spring constant on all C*α* atoms to reach a full dissociation. Observed initial distances of the centers-of-mass were equal to 28.70, 30.25, and 24.28A° for CRD2, CRD2e_K54E, and CRD2(39-73)e, respectively, indicating that the truncated variant formed a much more compact complex than the complete CRD2 molecules. It also suggests that the truncated amino-acid residues do not stick to the LIGHT or even prohibit other amino-acid residues from forming proper contacts. Although the CRD2e_K54E variant shows that larger work is needed for the dissociation, increased stability, compared to the wild-type, is observed only after about 3° A extension, and Fmax values of both variants are comparable (Fig. S10). The situation is completely different when CRD2(39-73)e is taken into consideration – it shows not only a significantly larger work needed for complete dissociation than both untruncated variants (24.68 ±6.16, 35.70 ±11.43, and 66.18 ±11.07 kcal/mol, respectively for CRD2, CRD2e_K54E, and CRD2(39-73)e) but also a much greater Fmax value is observed (1.409 ±0.795, 1.697 ±0.895, and 4.141 ±1.222kcal/mol/° A, respectively for CRD2, CRD2e_K54E, and CRD2(39-73)e). Moreover, truncation did not diminish the long-range interactions of the molecules; therefore, it should not have a negative impact on the recognition and early stages of the HVEM-fragment-LIGHT binding (Fig. S11).

### Designed truncated peptide is highly selective to the LIGHT binding pocket

As the LIGHT trimer consists of three identical binding sites, the analysis of the HDOCK results reveals that CRD2(39-73)e is docked to these three positions in its top 10 binding modes, with one of these modes ranking as ‘top1’. This finding suggests that CRD2(39-73)e and HVEM can effectively compete for the same binding site on the LIGHT trimer, therefore, inhibiting the HVEM-LIGHT complex formation. This, with the docking score of -342.41 and confidence score of 0.9791 shows strong preference of the CRD2(39-73)e peptide to bind to LIGHT trimer.

As this study primarily focuses on silencing the stimulatory signal with LIGHT in oppose to the inhibitory signal with BTLA, we also docked the best molecule candidates to BTLA. Docking of CRD2(39-73)e peptide to BTLA dimer shows low specificity, as the molecules in top10 binding modes are located in the interface between the molecules with the best docking score of -239.82 and confidence score of 0.8577, which are significantly worse than for the binding with LIGHT trimer. CRD2e_K54E and CRD2e show worse docking (-252.88 and -213.45, respectively) and confidence (0.8867 and 0.7806, respectively) scores when binding to the LIGHT, as well as lower selectivity comparing to binding to the BTLA molecule, making CRD2(39-73)e the best drug candidate. Moreover, calculated lRMSD values, representing the position of the CRD2e_K54E and CRD2(39-73)e in the LIGHT binding pocket was the lowest (most stable) among all tested variants (Fig. S12), further indicating high selectivity.

## Discussion

In this study, we investigated the stability of the HVEM molecule, its domains and fragments of these domains, and their interactions with the LIGHT trimer to uncover potential strategies for modulating LIGHT activity. Our extensive MD simulations of unbound HVEM and its variants, and in complex with LIGHT shed light on the role of disulfide bonds in domain stability, revealing their varying stabilizing properties. Although this is an understandable result, it is not always true that the addition of a disulfide bond increases protein stability. For example, in peroxiredoxin enzymes, disulfide bond formation introduces structural frustration and an increase in dynamics, which allows for a broader conformational landscape. (Troussicot *et al*., 2023).

Our MD simulations highlighted CRD2 as a hotspot for interactions with LIGHT, confirming experimental observations. Performed effective binding energy decomposition (Fig. S9) explains why experimental mutations of the selected amino-acid residues Y9, S20, Y23, R24, K26, E27, E38, R75 (Cheung *et al*., 2005), and D7, E14, S20, E31, L32 (Shrestha *et al*., 2020) do not affect the binding affinity to LIGHT, due to their near-zero contribution to LIGHT binding and presence in regions, which are not directly involved in binding. We found that L52 and M60 play an important role in the binding interface for the complete HVEM molecule (Fig. S9), while other experimentally determined amino-acid residues (Liu *et al*., 2021), such as H48 and L56, are in their vicinity an may play the role to stabilize the local structure of HVEM, rather than playing role in formation of the direct interactions. It is further confirmed by the observation of the Liu et al. that only pairwise mutagenesis of the H48, L52, and L56 has the observable effect on the LIGHT binding.

Work of the Shrestha et al., which shown that the G51F mutant of HVEM binds to LIGHT comparable to wt HVEM (Shrestha *et al*., 2021), is a good example that not all of the mutations of the amino-acid residues involved in the interaction with the partner molecule have the significant influence on the total binding affinity. In our study, we observed similar behaviour for some of the substitutions of the K54, C55, C61, D62, and N72, while some other amino-acid residues, such as P39, are much more sensitive for the substitution.

The presence of disulfide bonds emerged as a critical determinant of HVEM domain stability, with specific combinations exerting varying impacts on structural integrity. Our results suggest that these bonds play a pivotal role in maintaining the tertiary structure of HVEM domains, influencing their interactions with other proteins. Interestingly, we found that in some cases lack of disulfide bond allow structure to open-up and adjust during binding to LIGHT, for example CRD2no binds stronger that CRD2all, while increasing flexibility and binding strengh (Fig. S4,S5,S7). We successfully identified promising peptide inhibitors targeting the LIGHT trimer. Through MM-GBSA analyses, we discerned the CRD2 domain as the key binding site for LIGHT, offering potential for designing effective HVEM-based inhibitors. Mutational and truncational studies of the CRD2e peptide revealed valuable insights into enhancing binding affinity, albeit with intriguing complexities that warrant further investigation. Steered MD simulations provided dynamic insights into binding and dissociation events between HVEM fragments and LIGHT. The differential behavior of CRD2 variants during dissociation highlights the influence of truncation of specific amino-acid residues in stabilizing the complex, without impacting recognition of the molecules.

Overall, our study advances the understanding of the molecular basis of HVEM-LIGHT interactions and its domain stability, contributing to the development of targeted therapeutic interventions for immune-related disorders. The identified peptide inhibitors and the mechanistic insights gained from our simulations pave the way for future investigations in this direction, proposing a truncated variant of the CRD2, namely CRD2(39-73)e, as the most promising compound interacting selectively with LIGHT trimer. Moreover, our results can be used for further experimental verification as in our previous work.(Kuncewicz *et al*., 2023) Overall, the comprehensive multi-step study of structural properties, binding affinity, selectivity, and mechanisms establishes the gold standard for the computational design of new peptide drugs.

## Acknowledgment

This work has financial support from the National Science Centre, Poland, under SONATA No 2019/35/D/ST4/03156 (P.K. and P.S. - studies of disulfide-bond role) and PRELUDIUM Bis No 2020/39/O/ST4/01379 (P.C. and A.K.S. - studies of the LIGHT inhibitors).

