## Supporting file for "Computational Design of a Highly-Specific HVEM-Based Inhibitor of LIGHT Protein"

### Supporting Information for "Computational Design of a Highly-Specific HVEM-Based Inhibitor of LIGHT Protein"

#### 1 All-atom Molecular Dynamics simulations details

Molecular Dynamics (MD) simulations were performed using pmemd.cuda version 22 [1], which is a CUDA compatible GP-GPU variant of the pmemd [2], the main computational engine of the AMBER package [1]. Two main sets of MD simulations were run to analyze the influence of disulfide bonds on HVEM and its domains' stability with and without LIGHT: (i) without, and (ii) with the presence of LIGHT with different combinations of disulfide bonds in the HVEM molecule/fragment. The first set of MD simulations was used to determine the stability of the HVEM protein and its domains in an aquatic environment, while the second set was used to determine the binding affinity of proposed inhibitors to LIGHT. All MD simulations adapted the experimental model of HVEM-LIGHT (PDB: 4RSU, [3]) as the initial conformation, with or without disulfide-bond modifications, amino-acid substitutions and truncations. All systems were prepared using the tleap program with the ff19SB force field for proteins [4], OPC water model [5], and the co-optimized $Na^+$,$Cl^-$ ion models [6], as it is the recommended and most efficient combination of models and parameter sets for studies of protein-protein interactions using explicit water MD simulations in AMBER [1, 4].

In addition to classical MD simulations, a series of Steered MD simulations was run to establish a binding-unbinding mechanism for CRD2e, CRD2e_K54E, and CRD2(39-73)e representative models. To ensure that no inter-periodic contacts were formed during the simulation, a larger number of water molecules was added to form a layer of 30Å giving a truncated octahedron box dimensions of approximately 122x122x122Å. For those systems, 25 independent SMD trajectories were run with restraints placed on centers of mass of LIGHT and CRD2 molecules with force constant of 10 kcal/mol and pulling speed of 0.05 m/s to achieve full dissociation of the system (minimal distance of any heavy-atoms from the proteins above 6Å).

#### 2 Peptide synthesis

Selected peptides were synthesized by SPPS (Solid Phase Peptide Synthesis) method on LibertyBlue synthesizer using the Fmoc/tBu strategy. The resin used was Rink Amide Pro Tide (LL) with a capacity of 0.18 mmol/g. In order to selectively create disulfide bridges, two cysteine derivatives were used: Fmoc-Cys(Acm)-OH and Fmoc-Cys(Trt)-OH. Cysteine residues present in the amino acid sequence of HVEM and not involved in the disulfide bond were replaced with $\alpha$-aminobutyric acid (Abu), while methionine residues were replaced with norleucine (Nle) in order to avoid oxidation of sulfur in side chain. The reaction was carried out at room temperature for 24 hours. Then the peptides were cleavaged from the resin using a mixture consisting of 88% TFA, 5% phenol, 5% deionized water and 2% triisopropylsilane. 10 ml of the mixture was used per 1 g of resin and all were stirred for 4 hours at room temperature. Then resin was filtered off and $Et_2O$ was added to the remaining mixture to precipitate the peptide. The mixture was centrifuged three times at 4000 rpm for 15 minutes, at 4°C. Then the peptide was dissolved in deionized water and freeze-dried.

### 3 Peptide purification

The purification of the peptides was carried out on a reverse-phase high-performance liquid chromatography (RP-HPLC) on a Luna 5 $\mu$m C8(2) 100Å column. A 10-fold excess of DTT relative to free sulfhydryl groups was added to the aqueous solution of the purified peptide. The mixture was subjected to ultrasound at an 40°C temperature before being applied to the column. Two solutions were used during the purification: (A) - deionized water with 0.1% TFA (v/v), and (B) - 80% solution of acetonitrile in water with 0.08 % TFA (v/v). A linear concentration gradient from 5 to 50% B in A was applied over 120 minutes. The purification process was monitored using a UV detector measuring the absorption at 222 nm and 254 nm. Peptide purity was verified using RP-UHPLC with PDA and ELSD-LT detectors (SHIMADZU, Kyoto, Japan) Kromasil C8 analytical column Kinetex C8 (100 x 2.1 mm; 2.6 $\mu$m; 100Å) with using a 5-100% gradient of solution B in 15 minutes.

### 4 Disulfide bond formation

The first disulfide bond was formed between the cysteine residues with the sulfhydryl group protected by a trityl group, which was removed when the peptide was pulled from the resin. After purification, the peptide was dissolved in a mixture of water and methanol (1: 9, v/v) at a concentration of 40 mg/l. The pH of the mixture was adjusted to between 8 and 9 using ammonia water. The mixture was stirred for 7 days and compressed air run through the solution. The progress of the reaction was monitored by RP-HPLC. After this time, the methanol was evaporated and the remaining aqueous solution of the peptide was freeze-dried. The peptides with one disulfide bond was purified according to the procedure given earlier, while the peptides with two disulfides were subjected the second oxidation. The second disulfide bond was created according to the following procedure. The peptide was dissolved in a mixture of acetic acid, water and methanol (1:1:9, v/v/v) at a concentration of 40 mg/l. A 25-50 fold excess of iodine dissolved in methanol was then added to the mixture. The mixed solution was left for a week. After this time, the mixture was filtered through a Dowex ion exchange bed to remove iodine excess. The solvent was evaporated and the obtained peptide was dissolved in water and lyophilized. Then, the peptide was purified according to the procedure given earlier.

### 5 Influence of disulfide bonds on CRD stability

The HVEM domains demonstrate a very different role of particular disulfide bonds on their stability. In CRD1, out of all simulations with only one bond, Cys16-Cys29 (marked as b) lowered RMSD to 7.47Å from 11.20Å in simulations without bonds. Yet, only when coupled with Cys19-Cys29 (c), the RMSD lowers to 3.27Å, which is close to the result of simulations with all bonds present (2.89Å). Both the combinations "ab" and "ac" have results comparable to the one bond "b". It can be concluded that "a" has the smallest importance in the overall stability of CRD1.

In CRD3, which has up to two disulfide bonds, the presence of Cys83-Cys100 (g) has a greater effect on stability (3.89 Å) than the presence of Cys89-Cys97 (h). The measured end-to-end distance for CRD3h is comparable to that without bonds, and for CRD3g, it is comparable to all bonds present. The RMSF of residues in both the C- and N-terminal in simulations without (g) are much higher compared to those with (h) present (Fig. S5).

Further increase in the stability of the domain was observed when the rest of the HVEM was present, indicating that the these are rather protein fragments than structurally independent moieties.

The radius of gyration (RG) and RGmax analyses provide additional confirmation that, for CRD1, the most crucial bond is (b). This is evident from the similarity in values between simulations where it is present and those where all bonds are present, indicating a consistent level of compactness. In contrast, the compactness of CRD2 was not significantly affected by the presence or absence of disulfide bonds, except for CRD2no and CRD2d (Fig. S2), which exhibited a more relaxed or loose structure, in comparison to the structures with disulfide bonds in position (e) and (f). Regarding CRD3, disulfide bond (g) appears to contribute more significantly to structural stabilization than (h). It is worth noting that for molecules without disulfide bonds the rigidify occurs upon binding. The higher bonding affinity the more structure rigidity is observed (Fig S2, S3). The SASA of bound and unbound HVEM variants (Fig. S4) are similar (the variants bound to LIGHT trimer have slightly larger SASA), this indicate that hydrophobic interaction are not the key players in HVEM-LIGHT interactions.

S7 Number of native contacts between HVEM variant and LIGHT trimer as a function of time for last 20ns for (A): CRD1no (red), CRD1a (blue), and CRD1b (grey); (B): CRD1c (red), CRD1ab (blue), and CRD1ac (grey); (C): CRD1bc (red), CRD1abc (blue), and CRD1(16-38)no (grey); (D): CRD1(16-38)bc (red), CRD1-2(16-73)bce (blue), and CRD1-2(16-73)bce_K54E (grey) (E): CRD2no (red) CRD2no_K54E (blue), CRD2no_K54D (grey); (F): CRD2no_K54I (red); (F): CRD2no_K54L (blue), and CRD2no_K54S (grey); (G): CRD2no_K54V (red) CRD2no_K54Y (blue), and CRD2d (grey); (H): CRD2e (red), CRD2e_K54E (blue), and CRD2e_K54D (grey). Different trajectories are depicted as various color tone. . . . . . . . . . . . . . . . . . . . . . . . . . . . . . . . . . . . 16

S7 continued: (I): CRD2e_K54I (red), CRD2e_K54L (blue), and CRD2e_K54S (grey); (J) CRD2e_K54V (red), CRD2e_K54Y (blue), and CRD2e_K54E_D62A (grey); (K): CRD2e_K54E_D62K (red), CRD2e_K54E_D62L (blue), and CRD2e_K54E_D62S (grey); (L): CRD2f (red); CRD2de (blue) and CRD2de_K54E (grey); (M): CRD2de_K54D (red), CRD2de_K54I (blue), and CRD2de_K54L (grey); (N): CRD2de_K54S (red), CRD2df (blue), and CRD2ef (grey); (O): CRD2def (red), CRD2(39-73)e (blue), and CRD2e(39-73)_K54E (grey); (P): CRD2(39-73)_P39A (red), CRD2(39-73)e_P39V(blue), and CRD2(39-73)e_P39W (grey). Different trajectories are depicted as various color tone. . . . . . . . . 17

S7 continued (Q): CRD2(39-73)e_P39L(red), CRD2(39-73)e_P39I (blue), and CRD2(39-73)e_P39x (grey); (R): CRD2(39-73)e_N72A (red), CRD2(39-73)e_N72G(blue), and CRD2(39-73)e_C55A (grey); (S): CRD2(39-73)e_C55V(red), CRD2(39-73)e_C61S (blue), and CRD2(39-73)no (grey); (T): CRD3no (red), CRD3g (blue) and CRD2(39-73)h (grey) (S): CRD3gh (blue). Different trajectories are depicted as various color tone. . . . . . . . . 18

S8 Bar plot of the effective free energies of the various stable variants of the HVEM molecule interacting with the LIGHT trimer. . . . . . . . . 19

S9 MM-GBSA effective binding energy decomposition results for (A) CRD1no, (B) CRD1(16-38)no, (C) CRD1bc, (D) CRD1-2(16-73)bce, (E) CRD1-2(16-73)bce_K54E, (F) CRD2no variants . . . . . . . . . 20

S9 continued: MM-GBSA effective binding energy decomposition results for (G) CRD2no_K54D, (H) CRD2no_K54I, (I) CRD2no_K54L, (J) CRD2no_K54S, (K) CRD2no_K54V, (L) CRD2no_K54Y variants . . . . . . . . . 21

S9 continued: MM-GBSA effective binding energy decomposition results for: (M) CRD2d, (N) CRD2e, (O) CRD2e_K54D, (P) CRD2e_K54E, (Q) CRD2e_K54I, (R) CRD2e_K54L variants . . . . . . . . . 22

S9 continued: MM-GBSA effective binding energy decomposition results for: (S) CRD2e_K54S, (T) CRD2e_K54V, (U) CRD2e_K54Y, (V) CRD2e_K54E_D62A, (W) CRD2e_K54E_D62K, (X) CRD2e_K54E_D62L variants . . . . . . . . . 23

S9 continued: MM-GBSA effective binding energy decomposition results for: (Y) CRD2f, (Z) CRD2de, (AA) CRD2de_K54D, (AB) CRD2de_K54E, (AC) CRD2de_K54I, (AD) CRD2de_K54S variants . . . . . . . . . 24

S9 continued: MM-GBSA effective binding energy decomposition results for: (AE) CRD2df, (AF) CRD2ef, (AG) CRD2def, (AH) CRD2(39-73)no, (AI) CRD2(39-73)e, (AJ) CRD2(39-73)e_K54E variants . . . . . . . . . 25

S9 continued: MM-GBSA effective binding energy decomposition results for: (AK) CRD2(39-73)e_P39I, (AL) CRD2(39-73)e_P39L, (AM) CRD2(39-73)e_P39V, (AN) CRD2(39-73)e_P39W, (AO) CRD2(39-73)e_P39x, (AP) CRD2(39-73)e_K54E variants . . . . . . . . . 26

S9 continued: MM-GBSA effective binding energy decomposition results for: (AQ) CRD2(39-73)e_C55A, (AR) CRD2(39-73)e_C55V, (AS) CRD2(39-73)e_C61S, (AT) CRD2(39-73)e_N72A, (AU) CRD2(39-73)e_N72G, (AV) CRD3no variants . . . . 27

S9 continued: MM-GBSA effective binding energy decomposition results for: (AW) HVEMno variant, (AX) HVEMall . . . . . . . . . 28

S10 Plots of force and work averaged over 25 SMD trajectories (solid lines) with the corresponding standard deviation (SD) values (shown as semi-transparent areas) for HVEM variants interacting with the LIGHT trimer. Simulations were run until full dissociation was achieved. . . . . . . . . 29

S11 Contact heatmap as a function of residue index and extension (relative difference of the distance between centers of mass of HVEM variant and LIGHT trimer) averaged over 25 SMD simulations for HVEM variants with the LIGHT trimer (left panels: A, C, E), and LIGHT amino-acid residues with HVEM variants (right panels: B, D, F). The values are normalized in the range of 0-1. . . . . . . 30
S12 Bar plots of the strongly interacting complexes (ΔG>50kcal/mol) of the structural properties: (A) Complex C$\alpha$RMSD, (B) Receptor C$\alpha$RMSD, and (C) Ligand C$\alpha$RMSD values averaged for 3 trajectories for the last 20 ns of simulation. For the RMSD calculation, an initial conformation adapted from the PDB file was used as a reference. . . . . . . . . . . . . . . . . . . . . . . . . . . . . . . . 31
S12 continued: Bar plots of the strongly interacting complexes (ΔG>50kcal/mol) of the structural properties: (D) RG and (E) RGmax values averaged for 3 trajectories for the last 20 ns of simulation. For the RMSD calculation, an initial conformation adapted from the PDB file was used as a reference. . . . . . . . . . 32
S12 continued: Bar plots of the strongly interacting complexes (ΔG>50kcal/mol) of the structural properties: (F) End-to-end distance and (G) lRMSD values averaged for 3 trajectories for the last 20 ns of simulation. For the RMSD calculation, an initial conformation adapted from the PDB file was used as a reference. . . . 33

Table S1: Names and amino acid sequences of the designed peptides

| name | disulfide bonds | residues |
|---|---|---|
| CRD1no | without | L$^{1}$PSCKEDEYPV$^{11}$GSECCPKCSP$^{21}$GYRVKEACG$^{31}$ELTGTVCE |
| CRD1a | $C^{4}$-$C^{15}$ | L$^{1}$PS**C**KEDEYPV$^{11}$GSE**C**CPKCSP$^{21}$GYRVKEACG$^{31}$ELTGTVCE |
| CRD1b | $C^{16}$-$C^{29}$ | L$^{1}$PSCKEDEYPV$^{11}$GSEC**C**KCSP$^{21}$GYRVKEA**C**G$^{31}$ELTGTVCE |
| CRD1c | $C^{19}$-$C^{37}$ | L$^{1}$PSCKEDEYPV$^{11}$GSECCPK**C**SP$^{21}$GYRVKEACG$^{31}$ELTGTV**C**E |
| CRD1ab | $C^{4}$-$C^{15}$;$C^{16}$-$C^{29}$ | L$^{1}$PS**C**KEDEYPV$^{11}$GSE**CC**PKCSP$^{21}$GYRVKEA**C**G$^{31}$ELTGTVCE |
| CRD1ac | $C^{4}$-$C^{15}$;$C^{19}$-$C^{37}$ | L$^{1}$PS**C**KEDEYPV$^{11}$GSE**C**CPK**C**SP$^{21}$GYRVKEACG$^{31}$ELTGTV**C**E |
| CRD1bc | $C^{16}$-$C^{29}$;$C^{19}$-$C^{37}$ | L$^{1}$PSCKEDEYPV$^{11}$GSEC**C**PK**C**SP$^{21}$GYRVKEA**C**G$^{31}$ELTGTV**C**E |
| CRD1abc | $C^{4}$-$C^{15}$;$C^{16}$-$C^{29}$;$C^{19}$-$C^{37}$ | L$^{1}$PS**C**KEDEYPV$^{11}$GSE**CC**PK**C**SP$^{21}$GYRVKEA**C**G$^{31}$ELTGTV**C**E |
| CRD1(16-38)no | without | CPKCSP$^{21}$GYRVKEACG$^{31}$ELTGTVCE |
| CRD1(16-38)bc | $C^{16}$-$C^{29}$;$C^{19}$-$C^{37}$ | **C**PK**C**SP$^{21}$GYRVKEA**C**G$^{31}$ELTGTV**C**E |
| CRD1-2(16-73)bce | $C^{16}$-$C^{29}$;$C^{19}$-$C^{37}$;$C^{58}$-$C^{73}$ | **C**PK**C**SP$^{21}$GYRVKEA**C**G$^{31}$ELTGTV**C**EPCP$^{41}$PGTYIHLNG$^{51}$LSKCLQ**C**QMC$^{61}$DPAMGLRASR$^{71}$N**C**$^{73}$ |
| CRD1-2(16-73)bce_K54E | $C^{16}$-$C^{29}$;$C^{19}$-$C^{37}$;$C^{58}$-$C^{73}$ | **C**PK**C**SP$^{21}$GYRVKEA**C**G$^{31}$ELTGTV**C**EPCP$^{41}$PGTYIHLNG$^{51}$LSECLQ**C**QMC$^{61}$DPAMGLRASR$^{71}$N**C**$^{73}$ |
| CRD2no | without | PCP$^{41}$PGTYIHLNG$^{51}$LSKCLQCQMC$^{61}$DPAMGLRASR$^{71}$NCSRTENAVC$^{81}$ |
| CRD2no_K54D | without | PCP$^{41}$PGTYIHLNG$^{51}$LSDCLQCQMC$^{61}$DPAMGLRASR$^{71}$NCSRTENAVC$^{81}$ |
| CRD2no_K54E | without | PCP$^{41}$PGTYIHLNG$^{51}$LSECLQCQMC$^{61}$DPAMGLRASR$^{71}$NCSRTENAVC$^{81}$ |
| CRD2no_K54I | without | PCP$^{41}$PGTYIHLNG$^{51}$LSICLQCQMC$^{61}$DPAMGLRASR$^{71}$NCSRTENAVC$^{81}$ |
| CRD2no_K54L | without | PCP$^{41}$PGTYIHLNG$^{51}$LSLCLQCQMC$^{61}$DPAMGLRASR$^{71}$NCSRTENAVC$^{81}$ |
| CRD2no_K54S | without | PCP$^{41}$PGTYIHLNG$^{51}$LSSCLQCQMC$^{61}$DPAMGLRASR$^{71}$NCSRTENAVC$^{81}$ |
| CRD2no_K54V | without | PCP$^{41}$PGTYIHLNG$^{51}$LSVCLQCQMC$^{61}$DPAMGLRASR$^{71}$NCSRTENAVC$^{81}$ |
| CRD2no_K54Y | without | PCP$^{41}$PGTYIHLNG$^{51}$LSYCLQCQMC$^{61}$DPAMGLRASR$^{71}$NCSRTENAVC$^{81}$ |
| CRD2d | $C^{40}$-$C^{55}$ | P**C**P$^{41}$PGTYIHLNG$^{51}$LSK**C**LQCQMC$^{61}$DPAMGLRASR$^{71}$NCSRTENAVC$^{81}$ |
| CRD2e | $C^{58}$-$C^{73}$ | PCP$^{41}$PGTYIHLNG$^{51}$LSKCLQ**C**QMC$^{61}$DPAMGLRASR$^{71}$N**C**SRTENAVC$^{81}$ |
| CRD2e_K54D | $C^{58}$-$C^{73}$ | PCP$^{41}$PGTYIHLNG$^{51}$LSDCLQ**C**QMC$^{61}$DPAMGLRASR$^{71}$N**C**SRTENAVC$^{81}$ |
| CRD2e_K54E | $C^{58}$-$C^{73}$ | PCP$^{41}$PGTYIHLNG$^{51}$LSECLQ**C**QMC$^{61}$DPAMGLRASR$^{71}$N**C**SRTENAVC$^{81}$ |
| CRD2e_K54I | $C^{58}$-$C^{73}$ | PCP$^{41}$PGTYIHLNG$^{51}$LSICLQ**C**QMC$^{61}$DPAMGLRASR$^{71}$N**C**SRTENAVC$^{81}$ |
| CRD2e_K54L | $C^{58}$-$C^{73}$ | PCP$^{41}$PGTYIHLNG$^{51}$LSLCLQ**C**QMC$^{61}$DPAMGLRASR$^{71}$N**C**SRTENAVC$^{81}$ |
| CRD2e_K54S | $C^{58}$-$C^{73}$ | PCP$^{41}$PGTYIHLNG$^{51}$LSSCLQ**C**QMC$^{61}$DPAMGLRASR$^{71}$N**C**SRTENAVC$^{81}$ |
| CRD2e_K54V | $C^{58}$-$C^{73}$ | PCP$^{41}$PGTYIHLNG$^{51}$LSVCLQ**C**QMC$^{61}$DPAMGLRASR$^{71}$N**C**SRTENAVC$^{81}$ |
| CRD2e_K54Y | $C^{58}$-$C^{73}$ | PCP$^{41}$PGTYIHLNG$^{51}$LSYCLQ**C**QMC$^{61}$DPAMGLRASR$^{71}$N**C**SRTENAVC$^{81}$ |
| CRD2e_K54E_D62A | $C^{58}$-$C^{73}$ | PCP$^{41}$PGTYIHLNG$^{51}$LSECLQ**C**QMC$^{61}$APAMGLRASR$^{71}$N**C**SRTENAVC$^{81}$ |
| CRD2e_K54E_D62K | $C^{58}$-$C^{73}$ | PCP$^{41}$PGTYIHLNG$^{51}$LSECLQ**C**QMC$^{61}$KPAMGLRASR$^{71}$N**C**SRTENAVC$^{81}$ |
| CRD2e_K54E_D62L | $C^{58}$-$C^{73}$ | PCP$^{41}$PGTYIHLNG$^{51}$LSECLQ**C**QMC$^{61}$LPAMGLRASR$^{71}$N**C**SRTENAVC$^{81}$ |
| CRD2e_K54E_D62S | $C^{58}$-$C^{73}$ | PCP$^{41}$PGTYIHLNG$^{51}$LSECLQ**C**QMC$^{61}$SPAMGLRASR$^{71}$N**C**SRTENAVC$^{81}$ |
| CRD2f | $C^{61}$-$C^{81}$ | PCP$^{41}$PGTYIHLNG$^{51}$LSKCLQCQM**C**$^{61}$DPAMGLRASR$^{71}$NCSRTENAV**C**$^{81}$ |

Table S1: Continued: Names and amino acid sequences of the designed peptides (x denotes point deletion)

| | | |
|---|---|---|
| CRD2de | $C^{40}$-$C^{55}$;$C^{58}$-$C^{73}$ | P**C**P$^{41}$PGTYIHLNG$^{51}$LSK**C**LQ**C**QMC$^{61}$DPAMGLRASR$^{71}$N**C**SRTENAVC$^{81}$ |
| CRD2de_K54D | $C^{40}$-$C^{55}$;$C^{58}$-$C^{73}$ | P**C**P$^{41}$PGTYIHLNG$^{51}$LSD**C**LQ**C**QMC$^{61}$DPAMGLRASR$^{71}$N**C**SRTENAVC$^{81}$ |
| CRD2de_K54E | $C^{40}$-$C^{55}$;$C^{58}$-$C^{73}$ | P**C**P$^{41}$PGTYIHLNG$^{51}$LSE**C**LQ**C**QMC$^{61}$DPAMGLRASR$^{71}$N**C**SRTENAVC$^{81}$ |
| CRD2de_K54I | $C^{40}$-$C^{55}$;$C^{58}$-$C^{73}$ | P**C**P$^{41}$PGTYIHLNG$^{51}$LSI**C**LQ**C**QMC$^{61}$DPAMGLRASR$^{71}$N**C**SRTENAVC$^{81}$ |
| CRD2de_K54L | $C^{40}$-$C^{55}$;$C^{58}$-$C^{73}$ | P**C**P$^{41}$PGTYIHLNG$^{51}$LSL**C**LQ**C**QMC$^{61}$DPAMGLRASR$^{71}$N**C**SRTENAVC$^{81}$ |
| CRD2de_K54S | $C^{40}$-$C^{55}$;$C^{58}$-$C^{73}$ | P**C**P$^{41}$PGTYIHLNG$^{51}$LSS**C**LQ**C**QMC$^{61}$DPAMGLRASR$^{71}$N**C**SRTENAVC$^{81}$ |
| CRD2df | $C^{40}$-$C^{55}$;$C^{61}$-$C^{81}$ | P**C**P$^{41}$PGTYIHLNG$^{51}$LSK**C**LQCQM**C**$^{61}$DPAMGLRASR$^{71}$NCSRTENAV**C**$^{81}$ |
| CRD2ef | $C^{58}$-$C^{73}$;$C^{61}$-$C^{81}$ | PCP$^{41}$PGTYIHLNG$^{51}$LSKCLQ**C**QM**C**$^{61}$DPAMGLRASR$^{71}$N**C**SRTENAV**C**$^{81}$ |
| CRD2def | $C^{40}$-$C^{55}$;$C^{58}$-$C^{73}$;$C^{61}$-$C^{81}$ | P**C**P$^{41}$PGTYIHLNG$^{51}$LSK**C**LQ**C**QM**C**$^{61}$DPAMGLRASR$^{71}$N**C**SRTENAV**C**$^{81}$ |
| CRD2(39-73)e | $C^{58}$-$C^{73}$ | PCP$^{41}$PGTYIHLNG$^{51}$LSKCLQ**C**QMC$^{61}$DPAMGLRASR$^{71}$N**C**$^{73}$ |
| CRD2(39-73)e_P39A | $C^{58}$-$C^{73}$ | ACP$^{41}$PGTYIHLNG$^{51}$LSKCLQ**C**QMC$^{61}$DPAMGLRASR$^{71}$N**C**$^{73}$ |
| CRD2(39-73)e_P39L | $C^{58}$-$C^{73}$ | LCP$^{41}$PGTYIHLNG$^{51}$LSKCLQ**C**QMC$^{61}$DPAMGLRASR$^{71}$N**C**$^{73}$ |
| CRD2(39-73)e_P39I | $C^{58}$-$C^{73}$ | ICP$^{41}$PGTYIHLNG$^{51}$LSKCLQ**C**QMC$^{61}$DPAMGLRASR$^{71}$N**C**$^{73}$ |
| CRD2(39-73)e_P39V | $C^{58}$-$C^{73}$ | VCP$^{41}$PGTYIHLNG$^{51}$LSKCLQ**C**QMC$^{61}$DPAMGLRASR$^{71}$N**C**$^{73}$ |
| CRD2(39-73)e_P39W | $C^{58}$-$C^{73}$ | WCP$^{41}$PGTYIHLNG$^{51}$LSKCLQ**C**QMC$^{61}$DPAMGLRASR$^{71}$N**C**$^{73}$ |
| CRD2(39-73)e_P39x | $C^{57}$-$C^{72}$ | CP$^{40}$PGTYIHLNG$^{50}$LSKCLQ**C**QMC$^{60}$DPAMGLRASR$^{70}$N**C**$^{72}$ |
| CRD2(39-73)e_K54E | $C^{58}$-$C^{73}$ | PCP$^{41}$PGTYIHLNG$^{51}$LSECLQ**C**QMC$^{61}$DPAMGLRASR$^{71}$N**C**$^{73}$ |
| CRD2(39-73)e_C55A | $C^{58}$-$C^{73}$ | PCP$^{41}$PGTYIHLNG$^{51}$LSKALQ**C**QMC$^{61}$DPAMGLRASR$^{71}$N**C**$^{73}$ |
| CRD2(39-73)e_C55V | $C^{58}$-$C^{73}$ | PCP$^{41}$PGTYIHLNG$^{51}$LSKVLQ**C**QMC$^{61}$DPAMGLRASR$^{71}$N**C**$^{73}$ |
| CRD2(39-73)e_C61S | $C^{58}$-$C^{73}$ | PCP$^{41}$PGTYIHLNG$^{51}$LSKCLQ**C**QMS$^{61}$DPAMGLRASR$^{71}$N**C**$^{73}$ |
| CRD2(39-73)e_N72A | $C^{58}$-$C^{73}$ | PCP$^{41}$PGTYIHLNG$^{51}$LSKCLQ**C**QMC$^{61}$DPAMGLRASR$^{71}$A**C**$^{73}$ |
| CRD2(39-73)e_N72G | $C^{58}$-$C^{73}$ | PCP$^{41}$PGTYIHLNG$^{51}$LSKCLQ**C**QMC$^{61}$DPAMGLRASR$^{71}$G**C**$^{73}$ |
| CRD2(39-73)no | without | PCP$^{41}$PGTYIHLNG$^{51}$LSKCLQCQMS$^{61}$DPAMGLRASR$^{71}$NC$^{73}$ |
| CRD3no | without | GCSPQHFCIV$^{91}$QDGDHCAACR$^{101}$AYA |
| CRD3g | $C^{83}$-$C^{100}$ | G**C**SPQHFCIV$^{91}$QDGDHCAA**C**R$^{101}$AYA |
| CRD3h | $C^{89}$-$C^{97}$ | GCSPQHF**C**IV$^{91}$QDGDH**C**AACR$^{101}$AYA |
| CRD3gh | $C^{83}$-$C^{100}$;$C^{89}$-$C^{97}$ | G**C**SPQHF**C**IV$^{91}$QDGDH**C**AA**C**R$^{101}$AYA |

Table S2: Estimated secondary structure content

| | Estimated secondary structure content (%) | | | | | | | | | | | | | | | |
|---|---|---|---|---|---|---|---|---|---|---|---|---|---|---|---|---|
| | **$\alpha$-Helix** | | | | **$\beta$-sheet** | | | | **Turn** | | | | **Coil** | | | |
| **Peptide** | DSSP | STRIDE | KAKSI | CD | DSSP | STRIDE | KAKSI | CD | DSSP | STRIDE | KAKSI | CD | DSSP | STRIDE | KAKSI | CD |
| HVEM-CRD1no | 0.60 | 0.93 | | | 35.33 | 30.20 | | | 27.48 | 36.06 | | | 36.58 | 32.62 | | |
| CRD1no | 4.49 | 8.89 | 5.50 | | 10.26 | 12.47 | 9.20 | | 34.32 | 38.20 | | | 50.94 | 45.55 | 85.29 | |
| CRD1a | 4.58 | 10.82 | 7.77 | | 21.78 | 21.94 | 13.41 | | 30.28 | 35.22 | | | 43.36 | 37.60 | 78.82 | |
| CRD1b | 1.42 | 6.24 | 0.31 | | 20.55 | 16.22 | 8.23 | | 32.63 | 40.93 | | | 45.40 | 41.16 | 91.46 | |
| CRD1c | 2.45 | 5.64 | 0.35 | | 12.84 | 7.17 | 2.61 | | 33.80 | 43.39 | | | 50.91 | 49.60 | 97.04 | |
| CRD1ab | 3.02 | 6.18 | 0.73 | | 23.15 | 27.39 | 17.83 | | 30.98 | 34.84 | | | 42.85 | 36.36 | 81.44 | |
| CRD1ac | 1.90 | 6.49 | 1.87 | | 26.08 | 24.30 | 13.81 | | 31.42 | 42.30 | | | 40.59 | 30.75 | 84.32 | |
| CRD1bc | 0.54 | 3.39 | 0.57 | | 26.70 | 22.26 | 13.17 | | 27.87 | 41.68 | | | 44.89 | 33.75 | 86.26 | |
| CRD1abc | 0.02 | 3.07 | 0.00 | | 26.65 | 24.04 | 15.18 | | 27.66 | 42.82 | | | 45.66 | 32.84 | 84.82 | |
| HVEM-CRD1abc | 0.00 | 0.00 | | | 39.97 | 37.22 | | | 23.23 | 35.38 | | | 36.80 | 27.14 | | |
| HVEM-CRD2no | 1.87 | 3.07 | | | 12.39 | 10.89 | | | 29.86 | 31.55 | | | 55.87 | 52.28 | | |
| CRD2no | 5.22 | 8.13 | 6.46 | 3.9 | 10.44 | 4.48 | 0.34 | 28 | 32.44 | 39.50 | | 17.6 | 51.90 | 47.57 | 93.20 | 50.5 |
| CRD2d | 4.14 | 4.98 | 2.87 | 10.9 | 12.86 | 10.21 | 3.33 | 26.4 | 33.71 | 37.72 | | 15.4 | 49.30 | 46.38 | 93.81 | 47.3 |
| CRD2e | 2.58 | 0.89 | 0.09 | 13.3 | 20.28 | 17.48 | 16.21 | 18.8 | 28.65 | 37.92 | | 15 | 48.48 | 40.24 | 83.70 | 52.9 |
| CRD2e_K54E | 3.23 | 2.80 | 0.53 | 14.1 | 19.90 | 17.66 | 14.07 | 21.6 | 27.61 | 30.97 | | 15.1 | 49.26 | 46.70 | 85.40 | 49.2 |
| CRD2f | 2.84 | 3.44 | 3.76 | 7.9 | 13.28 | 13.71 | 11.27 | 23.3 | 31.97 | 40.68 | | 17.3 | 51.92 | 39.71 | 84.97 | 51.5 |
| CRD2de | 1.56 | 0.32 | 0.02 | 3.2 | 19.48 | 18.07 | 8.88 | 32.1 | 30.96 | 37.65 | | 17.4 | 48.00 | 42.97 | 91.10 | 47.3 |
| CRD2df | 1.38 | 2.95 | 0.00 | 1.2 | 20.84 | 20.34 | 7.44 | 39.5 | 34.66 | 42.10 | | 15.1 | 43.12 | 33.37 | 92.56 | 44.2 |
| CRD2ef | 1.32 | 1.08 | 0.10 | 2.5 | 19.63 | 19.77 | 16.11 | 33.4 | 28.15 | 32.18 | | 17.4 | 50.90 | 44.84 | 83.79 | 46.7 |
| CRD2def | 1.72 | 0.87 | 0.00 | | 19.67 | 18.46 | 9.88 | | 31.86 | 39.83 | | | 46.75 | 38.18 | 90.12 | |
| CRD2(39-73)e | 6.61 | 9.42 | 11.10 | | 9.09 | 2.13 | 5.67 | | 32.29 | 28.69 | | | 52.10 | 88.44 | 54.37 | |
| HVEM-CRD2def | 5.09 | 6.01 | | | 17.45 | 13.72 | | | 24.77 | 25.31 | | | 52.69 | 50.56 | | |
| HVEM-CRD3no | 2.31 | 4.29 | | | 20.03 | 8.55 | | | 31.66 | 37.64 | | | 46.00 | 49.32 | | |
| CRD3no | 7.56 | 21.78 | 13.58 | | 9.44 | 3.55 | 0.05 | | 33.10 | 35.75 | | | 49.90 | 54.49 | 86.37 | |
| CRD3g | 6.31 | 14.76 | 0.00 | | 29.96 | 27.90 | 16.19 | | 35.73 | 43.12 | | | 28.00 | 40.41 | 83.81 | |
| CRD3h | 1.45 | 14.38 | 1.13 | | 24.19 | 21.23 | 0.24 | | 30.29 | 32.36 | | | 44.06 | 54.11 | 98.63 | |
| CRD3gh | 0.03 | 14.60 | 0.00 | | 31.88 | 38.86 | 18.22 | | 28.17 | 35.28 | | | 39.92 | 37.32 | 81.78 | |
| HVEM-CRD3gh | 0.00 | 0.00 | | | 46.69 | 50.89 | | | 24.62 | 24.24 | | | 28.68 | 24.66 | | |

Table S3: Solvent accessible surfaces area (SASA) [Å$^2$] obtained for last 20 ns averaged over three trajectories and their standard deviation between trajectories.

| Name | SASA of complex | Sum of SASA of components | Ratio |
|---|---|---|---|
| CRD1no | 21793 ± 123 | 22566 ± 259 | 0.966 ± 0.006 |
| CRD1bc | 25262 ± 2090 | 25763 ± 1755 | 0.980 ± 0.015 |
| CRD1(16-38)no | 23833 ± 1703 | 24237 ± 2002 | 0.984 ± 0.012 |
| CRD1-2(16-73)bce | 25231 ± 1528 | 25737 ± 1404 | 0.980 ± 0.016 |
| CRD1-2(16-73)bce_K54E | 24095 ± 578 | 25235 ± 578 | 0.955 ± 0.024 |
| CRD2no | 25123 ± 2189 | 25582 ± 1986 | 0.982 ± 0.023 |
| CRD2no_K54D | 24743 ± 1492 | 25361 ± 1314 | 0.975 ± 0.018 |
| CRD2no_K54I | 24971 ± 2016 | 25456 ± 1592 | 0.980 ± 0.028 |
| CRD2no_K54L | 21883 ± 403 | 23306 ± 445 | 0.939 ± 0.002 |
| CRD2no_K54S | 24718 ± 2741 | 25081 ± 2421 | 0.984 ± 0.015 |
| CRD2no_K54V | 24767 ± 2205 | 25474 ± 1805 | 0.971 ± 0.021 |
| CRD2no_K54Y | 26345 ± 2527 | 26944 ± 2218 | 0.977 ± 0.016 |
| CRD2def | 24550 ± 1650 | 25751 ± 1548 | 0.953 ± 0.007 |
| CRD2d | 23337 ± 1531 | 24674 ± 1702 | 0.946 ± 0.004 |
| CRD2e | 24237 ± 2004 | 24922 ± 1860 | 0.972 ± 0.024 |
| CRD2e_K54D | 23486 ± 324 | 23842 ± 439 | 0.985 ± 0.007 |
| CRD2e_K54E | 24103 ± 1751 | 24662 ± 1499 | 0.977 ± 0.025 |
| CRD2e_K54I | 25385 ± 1477 | 26133 ± 1708 | 0.972 ± 0.007 |
| CRD2e_K54L | 22874 ± 1091 | 24023 ± 554 | 0.952 ± 0.037 |
| CRD2e_K54S | 24145 ± 1782 | 24429 ± 1542 | 0.988 ± 0.017 |
| CRD2e_K54V | 23837 ± 1628 | 24581 ± 1306 | 0.969 ± 0.016 |
| CRD2e_K54Y | 23456 ± 3045 | 24879 ± 2996 | 0.942 ± 0.011 |
| CRD2e_K54E_D62A | 22354 ± 595 | 23329 ± 557 | 0.959 ± 0.030 |
| CRD2e_K54E_D62K | 25340 ± 2252 | 25662 ± 2122 | 0.987 ± 0.012 |
| CRD2e_K54E_D62L | 23559 ± 608 | 24065 ± 750 | 0.979 ± 0.019 |
| CRD2f | 24639 ± 1542 | 25361 ± 1181 | 0.971 ± 0.021 |
| CRD2de | 23300 ± 1938 | 24183 ± 1870 | 0.963 ± 0.019 |
| CRD2de_K54D | 22298 ± 949 | 22956 ± 719 | 0.971 ± 0.012 |
| CRD2de_K54S | 21582 ± 744 | 22400 ± 501 | 0.963 ± 0.018 |
| CRD2de_K54E | 22901 ± 464 | 24198 ± 594 | 0.947 ± 0.013 |
| CRD2de_K54I | 22798 ± 376 | 23902 ± 88 | 0.954 ± 0.018 |
| CRD2df | 21272 ± 165 | 22639 ± 254 | 0.940 ± 0.004 |
| CRD2ef | 24146 ± 1967 | 25097 ± 2028 | 0.962 ± 0.017 |
| CRD2(39-73)no | 22343 ± 209 | 23489 ± 492 | 0.951 ± 0.012 |
| CRD2(39-73)e | 23765 ± 2148 | 24621 ± 1688 | 0.964 ± 0.034 |
| CRD2(39-73)e_P39A | 23400 ± 2209 | 23953 ± 2368 | 0.977 ± 0.017 |
| CRD2(39-73)e_P39I | 24892 ± 1753 | 25883 ± 1624 | 0.961 ± 0.011 |
| CRD2(39-73)e_P39L | 22699 ± 133 | 23398 ± 57 | 0.970 ± 0.004 |
| CRD2(39-73)e_P39V | 25384 ± 1813 | 25852 ± 2016 | 0.983 ± 0.015 |
| CRD2(39-73)e_P39W | 23523 ± 1889 | 24169 ± 2042 | 0.974 ± 0.019 |
| CRD2(39-73)e_P39x | 24127 ± 1826 | 24842 ± 1282 | 0.970 ± 0.024 |
| CRD2(39-73)e_C55A | 24474 ± 1943 | 25498 ± 1605 | 0.959 ± 0.022 |
| CRD2(39-73)e_C55V | 23251 ± 2839 | 24533 ± 2756 | 0.947 ± 0.009 |
| CRD2(39-73)e_C61S | 23123 ± 365 | 24232 ± 557 | 0.954 ± 0.015 |
| CRD2(39-73)e_N72A | 23896 ± 2472 | 24779 ± 2119 | 0.963 ± 0.033 |
| CRD2(39-73)e_N72G | 22300 ± 454 | 23785 ± 470 | 0.938 ± 0.006 |
| CRD3no | 24141 ± 849 | 24272 ± 857 | 0.995 ± 0.005 |
| HVEMno | 26750 ± 223 | 28726 ± 229 | 0.931 ± 0.011 |
| HVEMall | 25892 ± 2028 | 27787 ± 1332 | 0.931 ± 0.030 |

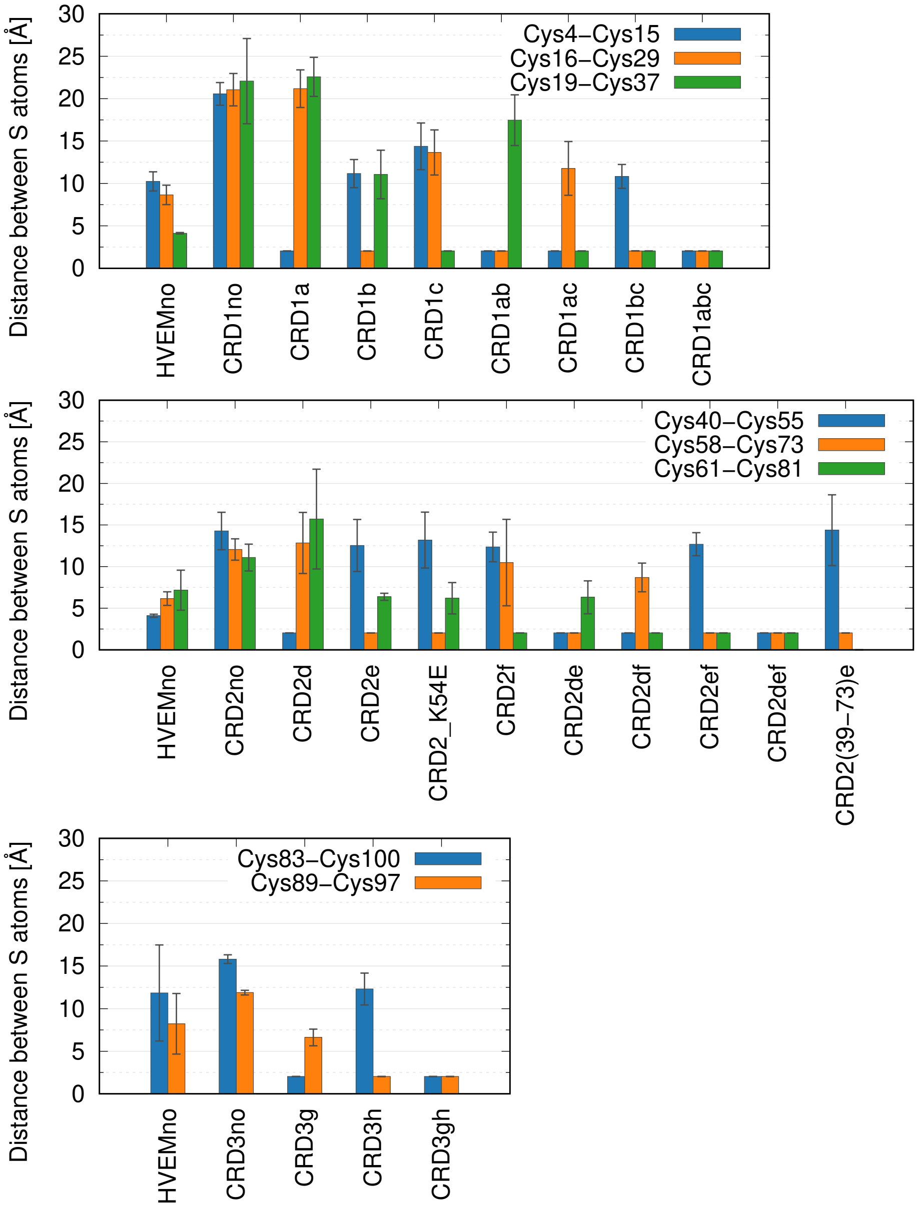

Figure S1: Distance between S atoms in disulfide bonds averaged for 3 trajectories for the second halves of the simulations (500-1000ns) for CRD1, CRD2 and CRD3 without the LIGHT trimer.

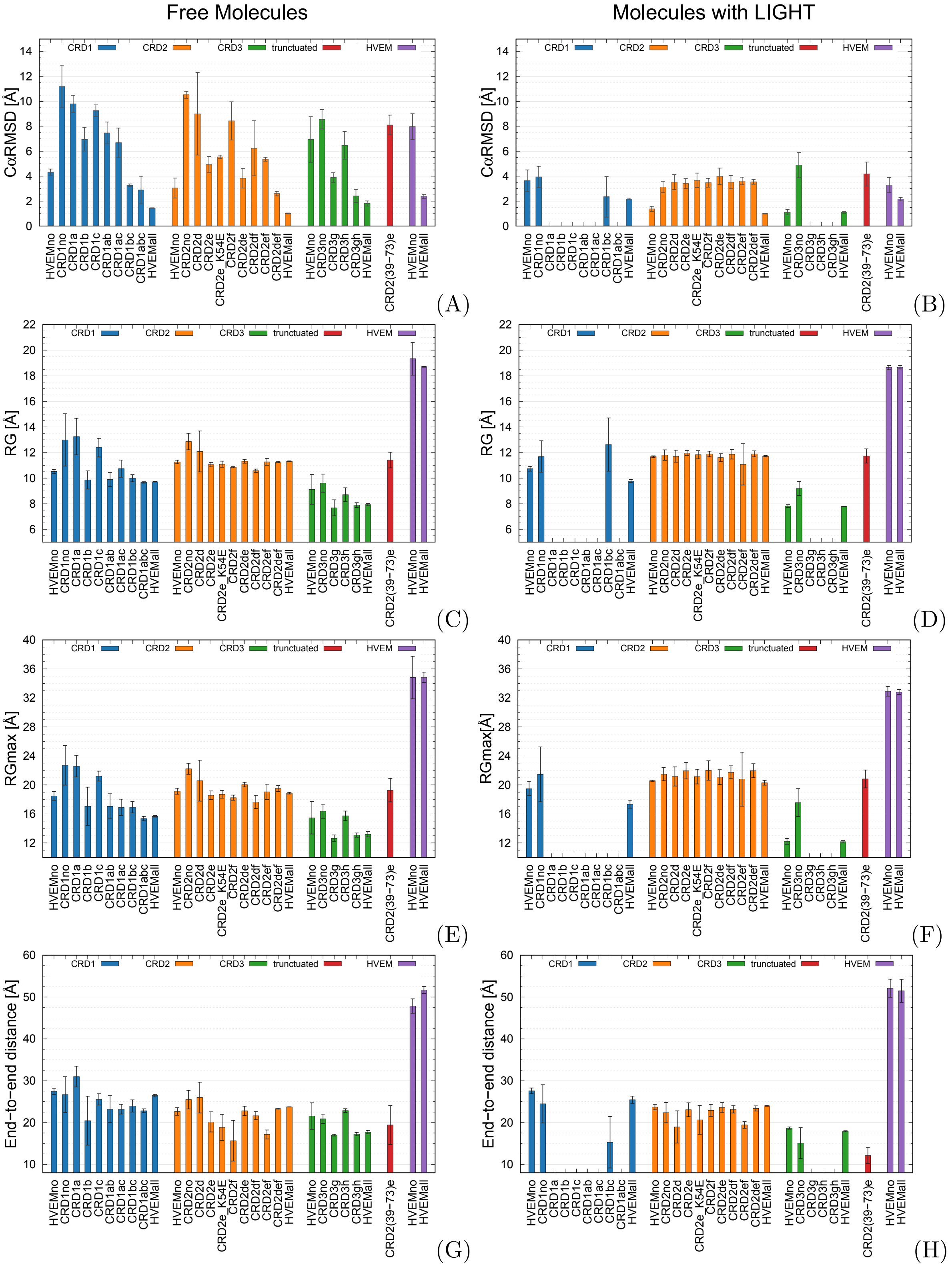

Figure S2: (A) RMSD, (C) RG, (E) RGmax, (G) End-to-end distance calculated from simulations of free domains and (B) RMSD, (D) RG, (F) RGmax and (H) End-to-end distance values calculated from simulations with LIGHT, averaged over the second halves of the 3 trajectories.

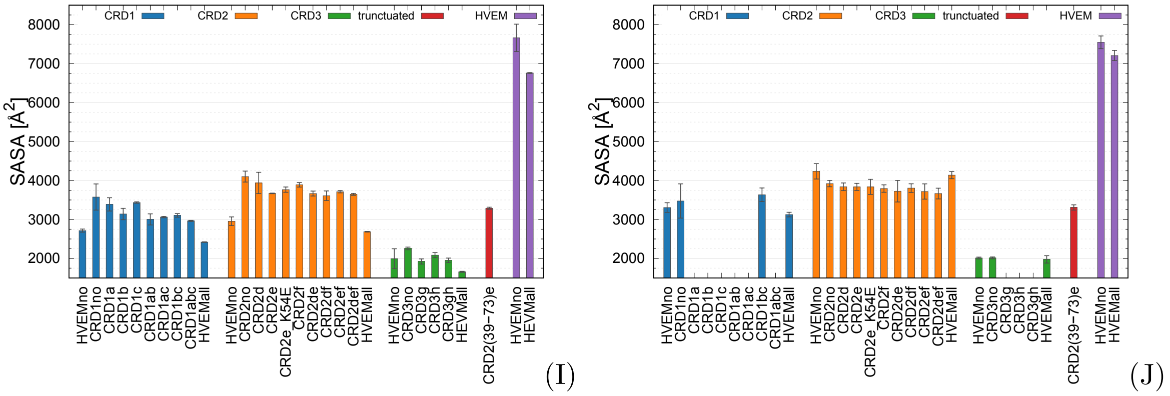

Figure S2: continued: (I) SASA values calculated from simulation of free domains and (J) theoretical SASA values calculated from simulation with the LIGHT trimer (the LIGHT trimer was excluded in this analysis, therefore, the whole HVEM variant surface was computed) averaged over the second halves of the 3 trajectories.

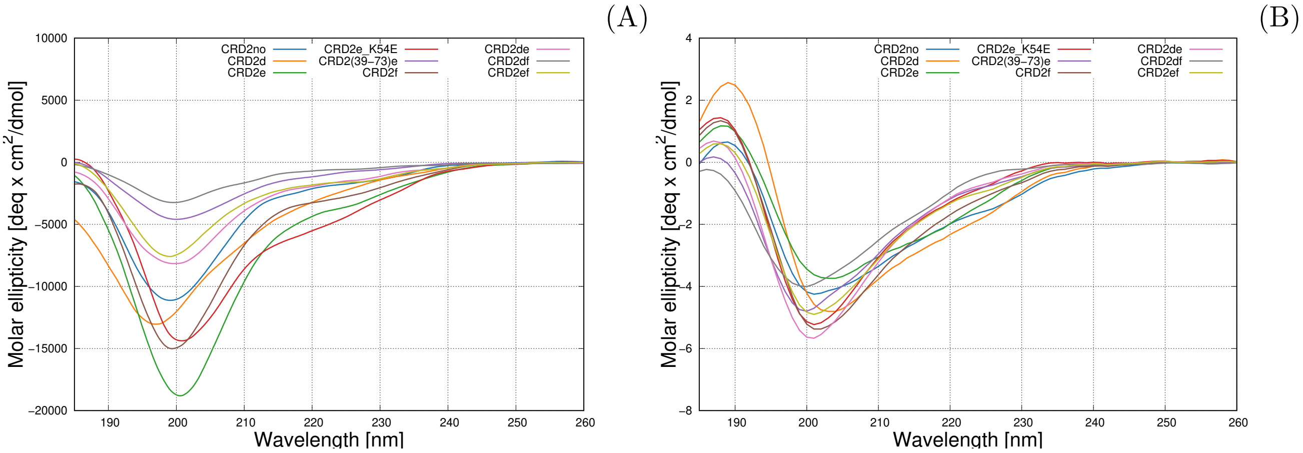

Figure S3: Experimental (A) and simulated (B) CD spectra of the selected designed peptides.

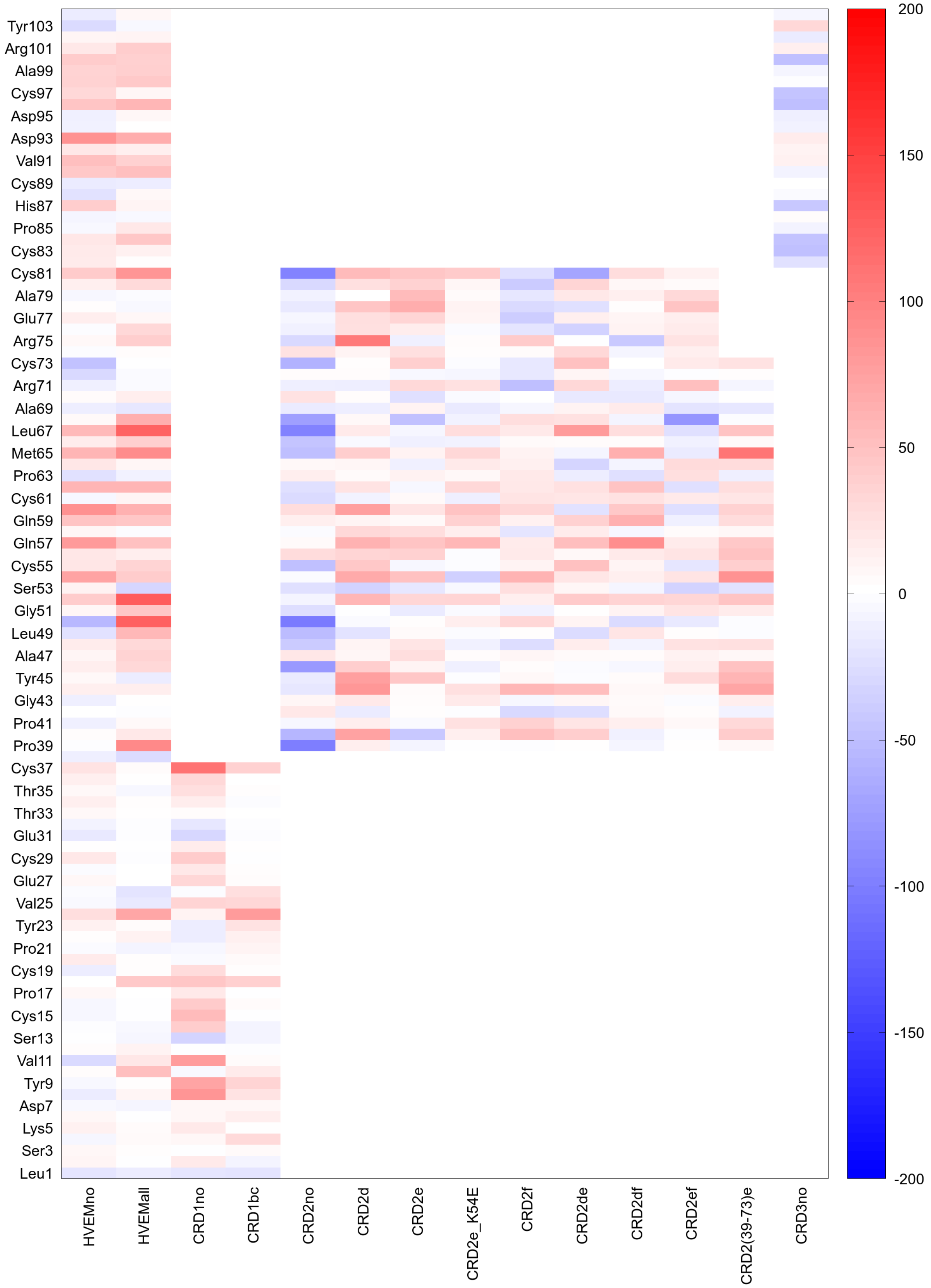

Figure S4: Heatmap of per-residue $\Delta SASA = SASA_{\text{free}} - SASA_{\text{complex}}$ calculated for all combinations of disulfide bonds present and absent in the simulations for the full HVEM molecule, its domains (CRD1-3), and fragment (CRD(39-73)e), averaged over the second halves of the 3 trajectories.

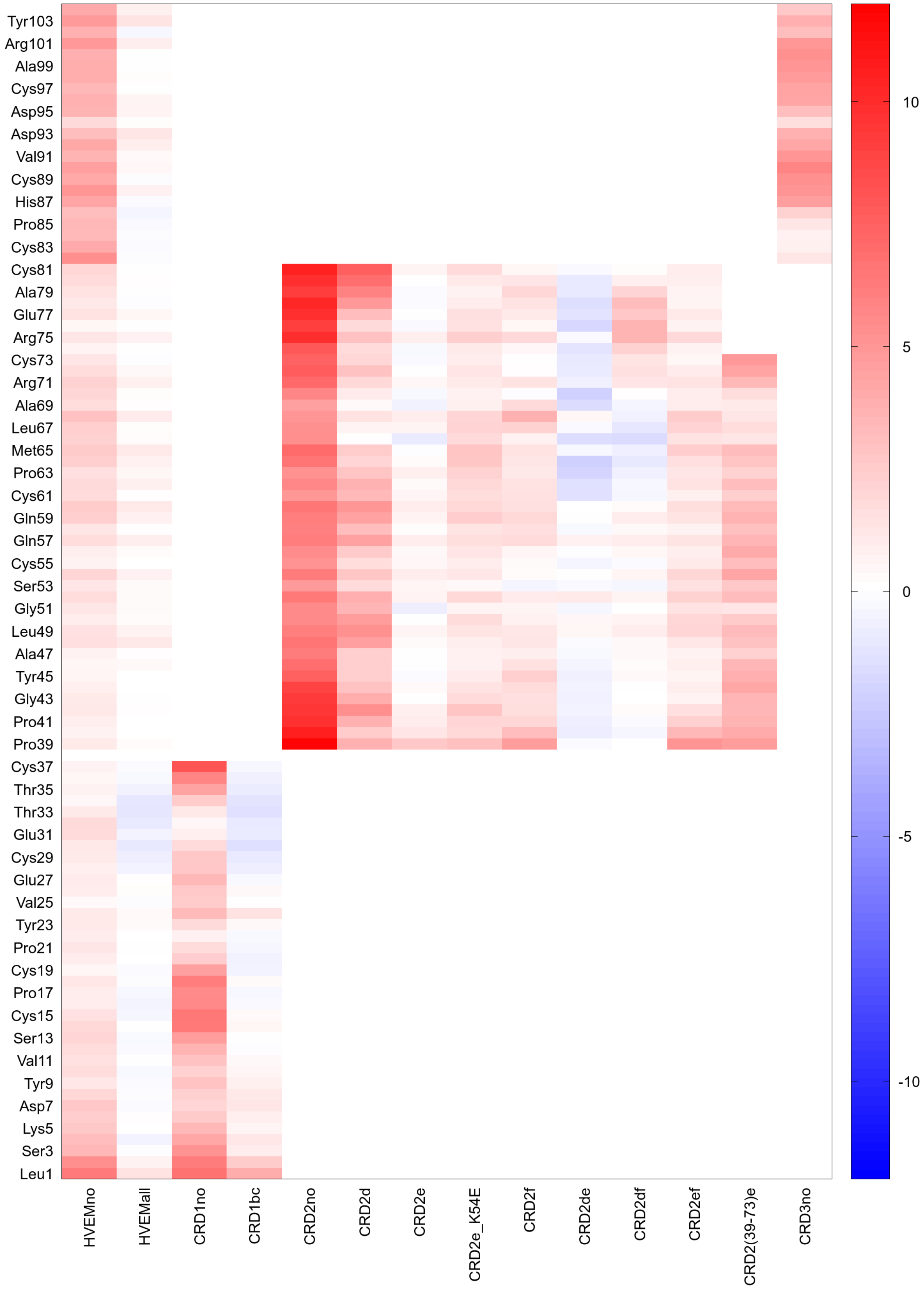

Figure S5: Heatmap of per-residue $\Delta RMSF = RMSF_{\text{free}} - RMSF_{\text{complex}}$ calculated for HVEM with and without disulfide bonds, and its selected variants forming stable complexes with LIGHT, averaged over the second halves of the 3 trajectories.

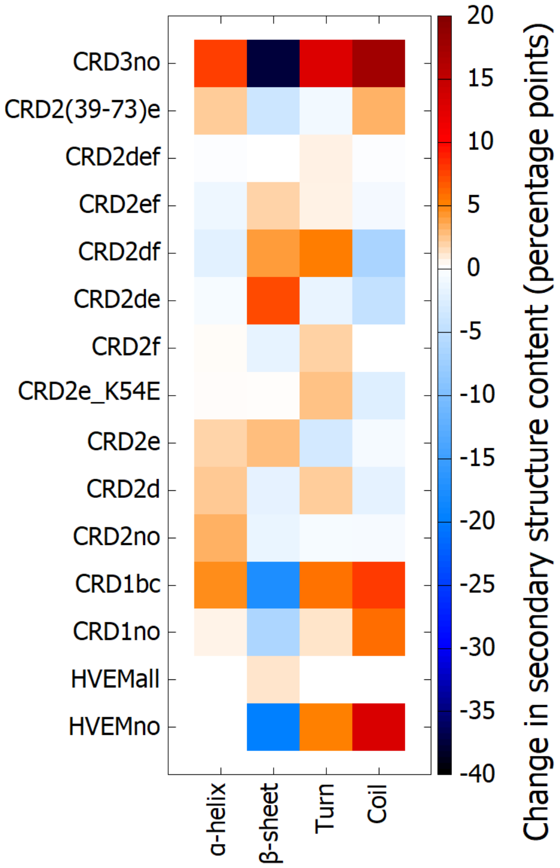

Figure S6: Heatmap of the change in the content of secondary structures expressed in percentage points calculated for HVEM with and without disulfide bonds, and its selected variants forming stable complexes with LIGHT, averaged over the second halves of the 3 trajectories

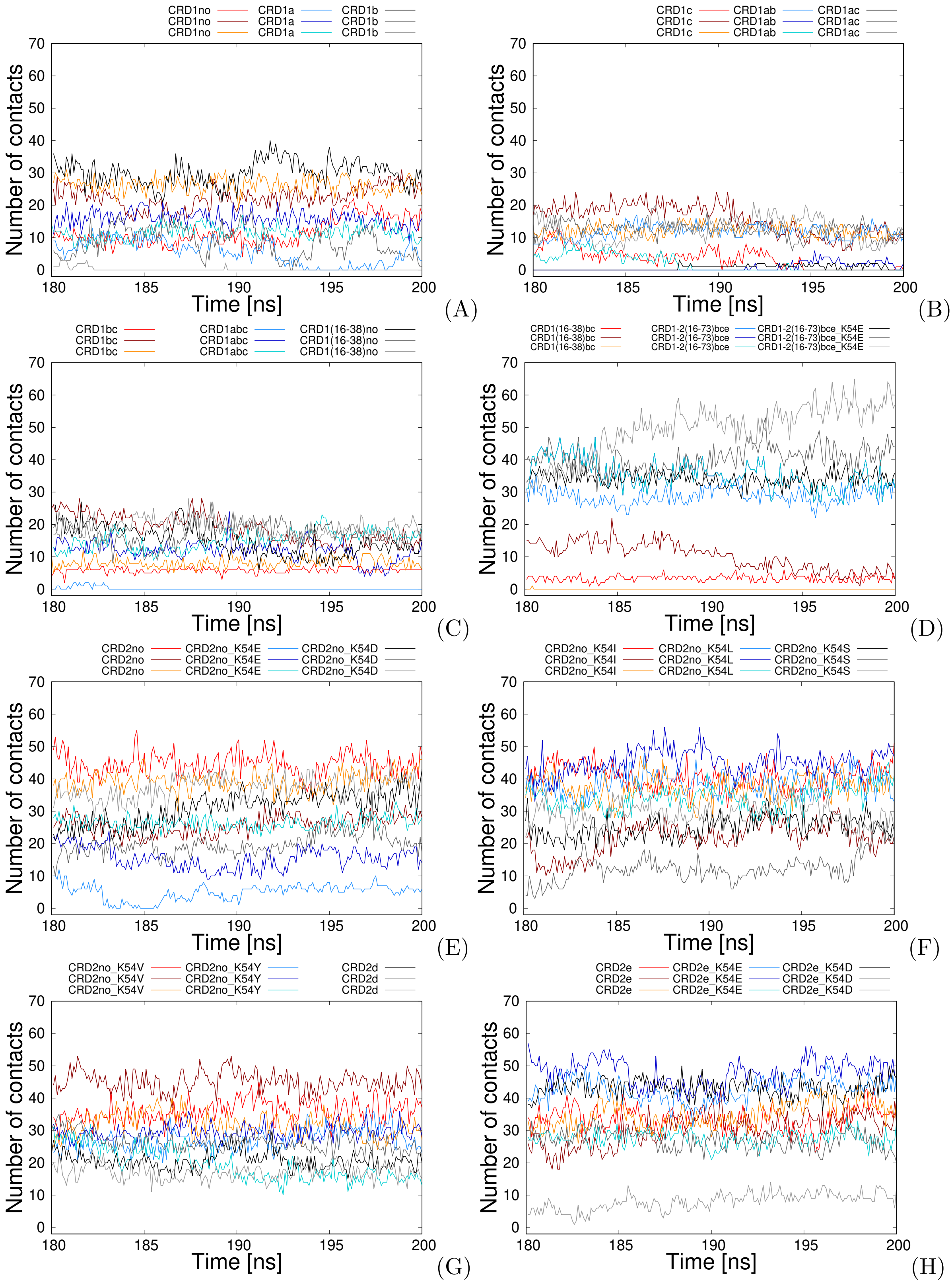

Figure S7: Number of native contacts between HVEM variant and LIGHT trimer as a function of time for last 20ns for (A): CRD1no (red), CRD1a (blue), and CRD1b (grey); (B): CRD1c (red), CRD1ab (blue), and CRD1ac (grey); (C): CRD1bc (red), CRD1abc (blue), and CRD1(16-38)no (grey); (D): CRD1(16-38)bc (red), CRD1-2(16-73)bce (blue), and CRD1-2(16-73)bce_K54E (grey) (E): CRD2no (red) CRD2no_K54E (blue), CRD2no_K54D (grey); (F): CRD2no_K54I (red); (F): CRD2no_K54L (blue), and CRD2no_K54S (grey); (G): CRD2no_K54V (red) CRD2no_K54Y (blue), and CRD2d (grey); (H): CRD2e (red), CRD2e_K54E (blue), and CRD2e_K54D (grey). Different trajectories are depicted as various color tone.

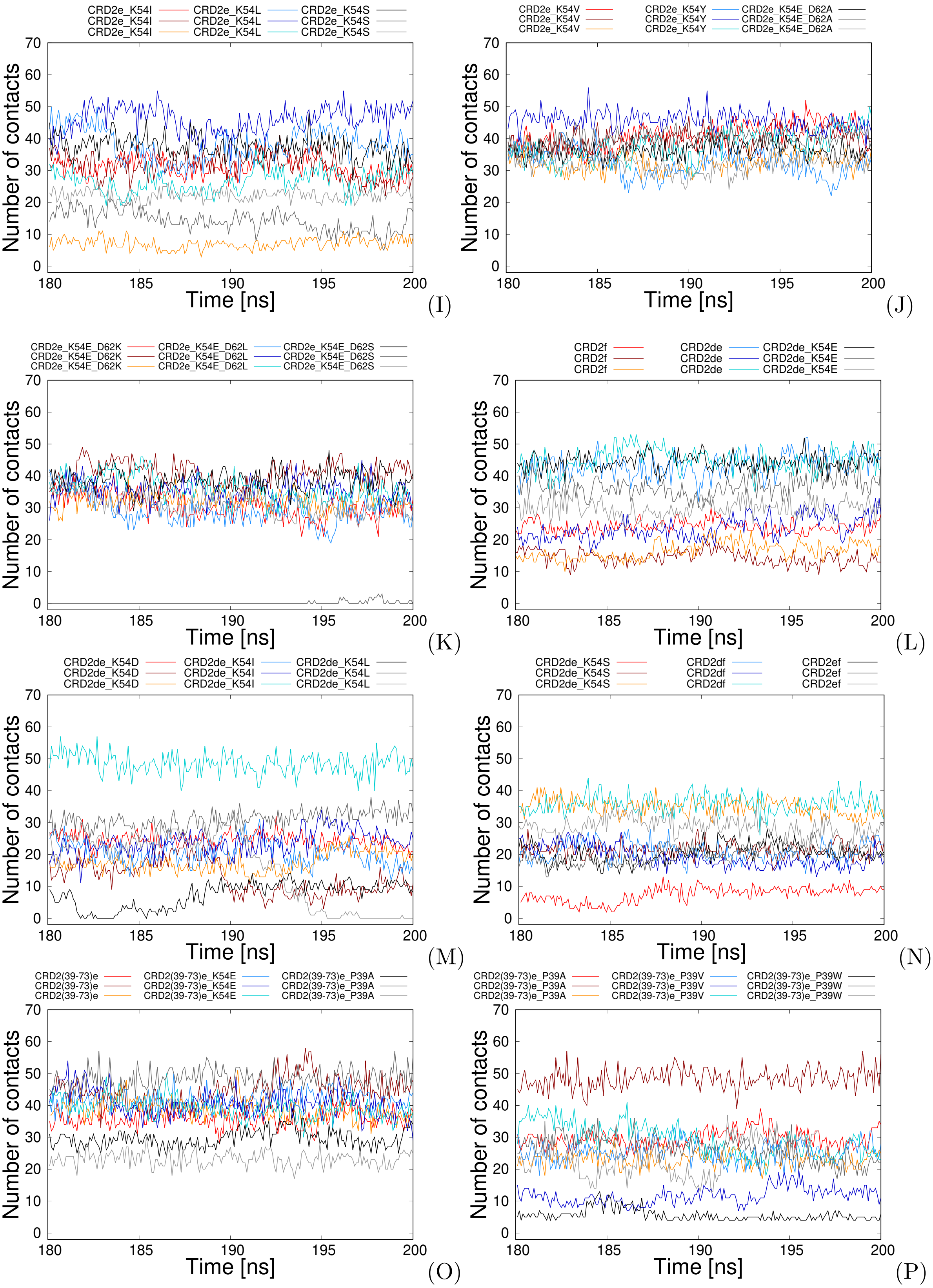

Figure S7: continued: (I): CRD2e_K54I (red), CRD2e_K54L (blue), and CRD2e_K54S (grey); (J) CRD2e_K54V (red), CRD2e_K54Y (blue), and CRD2e_K54E_D62A (grey); (K): CRD2e_K54E_D62K (red), CRD2e_K54E_D62L (blue), and CRD2e_K54E_D62S (grey); (L): CRD2f (red); CRD2de (blue) and CRD2de_K54E (grey); (M): CRD2de_K54D (red), CRD2de_K54I (blue), and CRD2de_K54L (grey); (N): CRD2de_K54S (red), CRD2df (blue), and CRD2ef (grey); (O): CRD2def (red), CRD2(39-73)e (blue), and CRD2e(39-73)_K54E (grey); (P): CRD2(39-73)_P39A (red), CRD2(39-73)e_P39V(blue), and CRD2(39-73)e_P39W (grey). Different trajectories are depicted as various color tone.

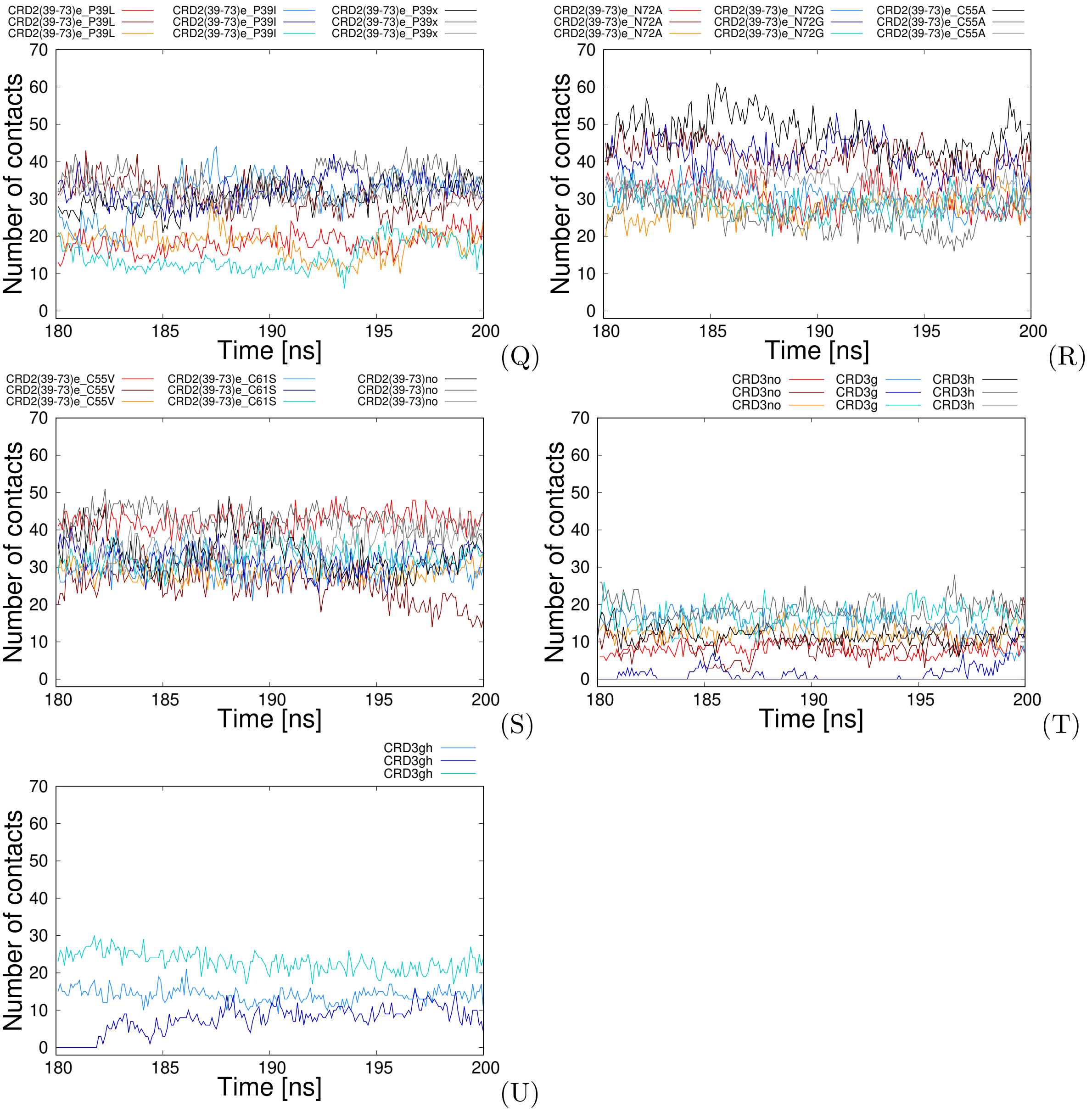

Figure S7: continued (Q): CRD2(39-73)e_P39L(red), CRD2(39-73)e_P39I (blue), and CRD2(39-73)e_P39x (grey); (R): CRD2(39-73)e_N72A (red), CRD2(39-73)e_N72G(blue), and CRD2(39-73)e_C55A (grey); (S): CRD2(39-73)e_C55V(red), CRD2(39-73)e_C61S (blue), and CRD2(39-73)no (grey); (T): CRD3no (red), CRD3g (blue) and CRD2(39-73)h (grey) (S): CRD3gh (blue). Different trajectories are depicted as various color tone.

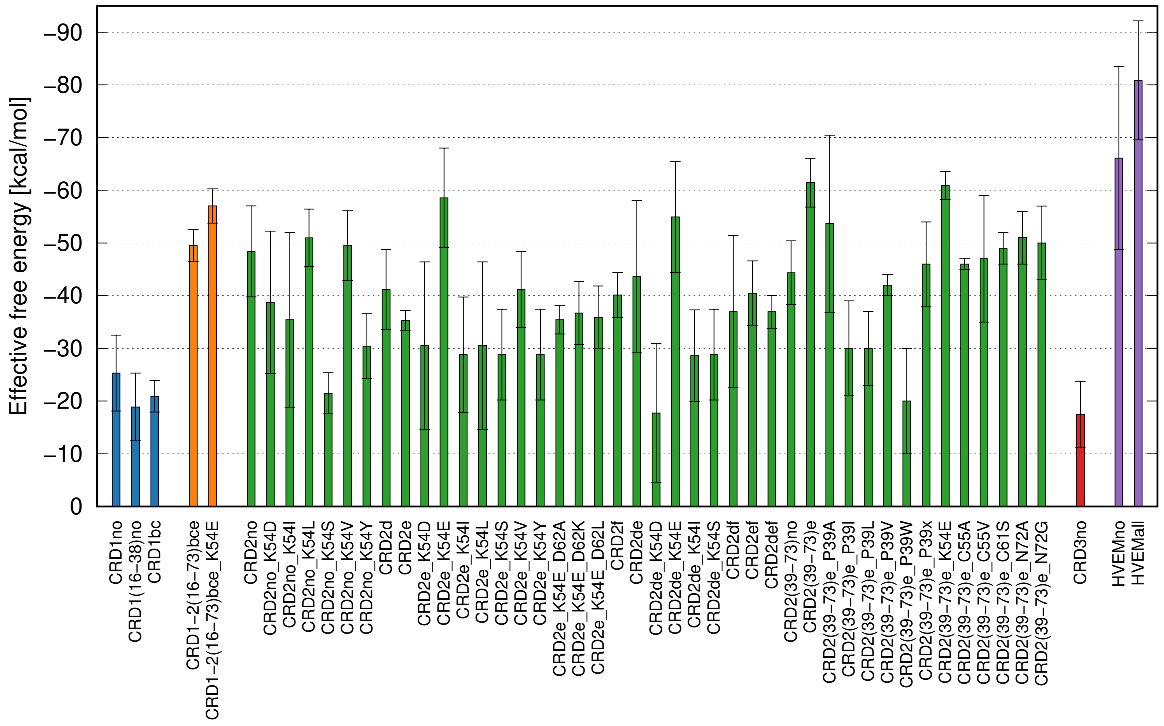

Figure S8: Bar plot of the effective free energies of the various stable variants of the HVEM molecule interacting with the LIGHT trimer.

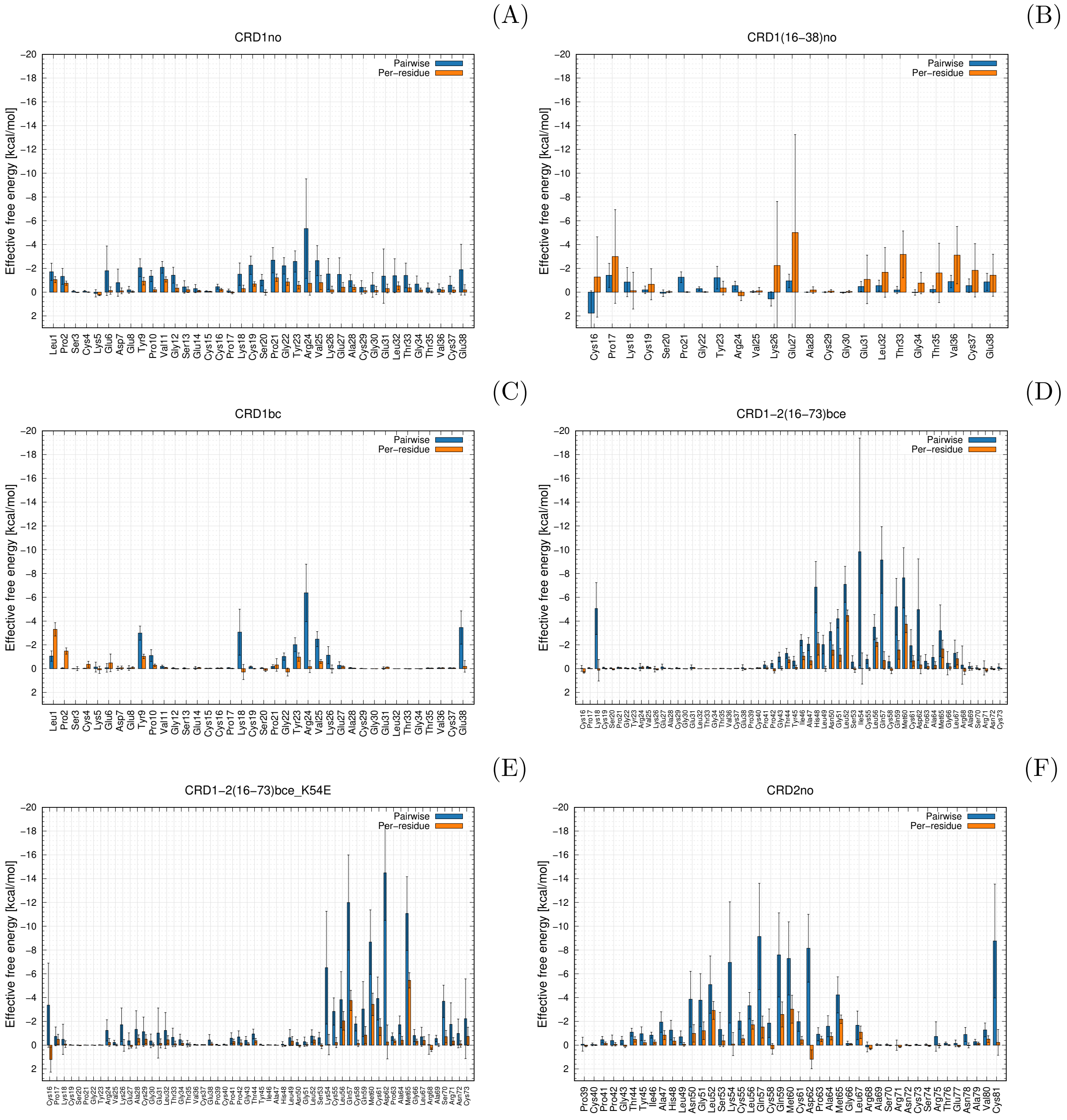

Figure S9: MM-GBSA effective binding energy decomposition results for (A) CRD1no, (B) CRD1(16-38)no, (C) CRD1bc, (D) CRD1-2(16-73)bce, (E) CRD1-2(16-73)bce_K54E, (F) CRD2no variants

(G)

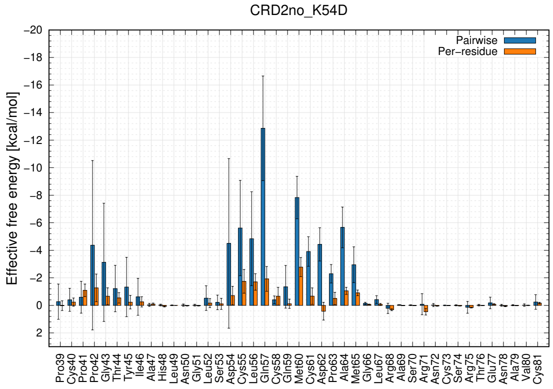

(H)

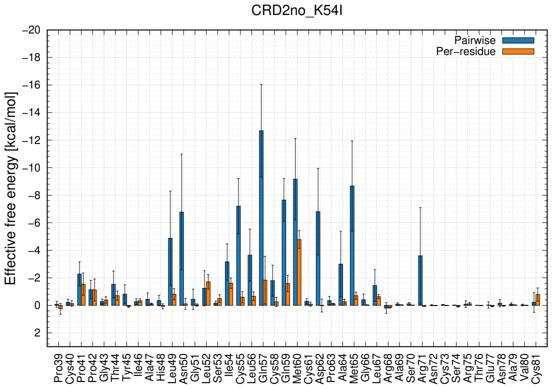

(I)

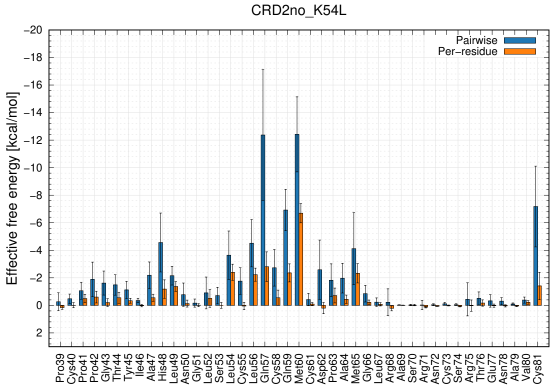

(J)

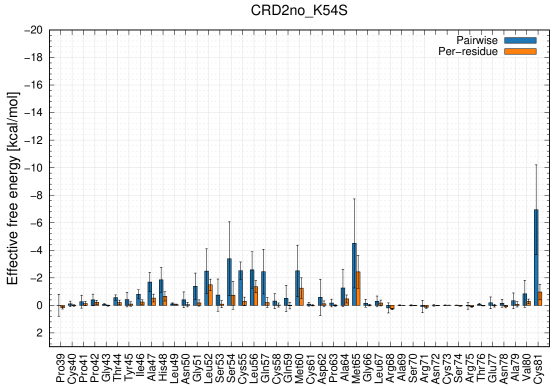

(K)

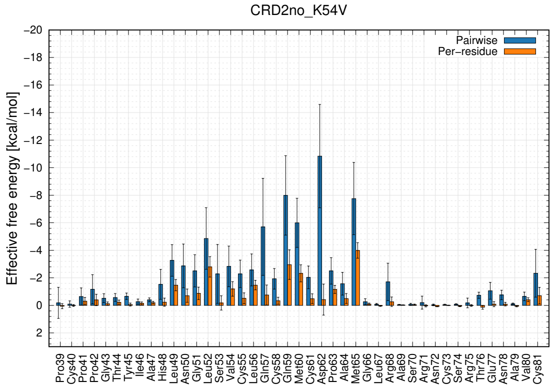

(L)

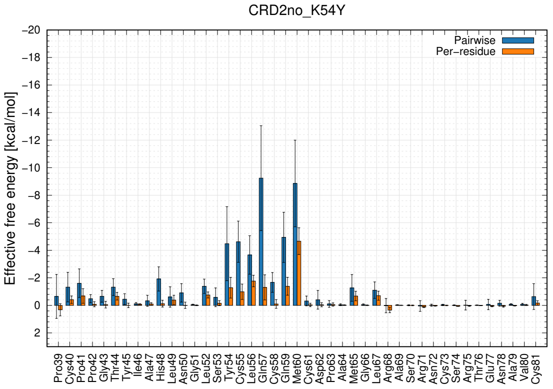

Figure S9: continued: MM-GBSA effective binding energy decomposition results for (G) CRD2no_K54D, (H) CRD2no_K54I, (I) CRD2no_K54L, (J) CRD2no_K54S, (K) CRD2no_K54V, (L) CRD2no_K54Y variants

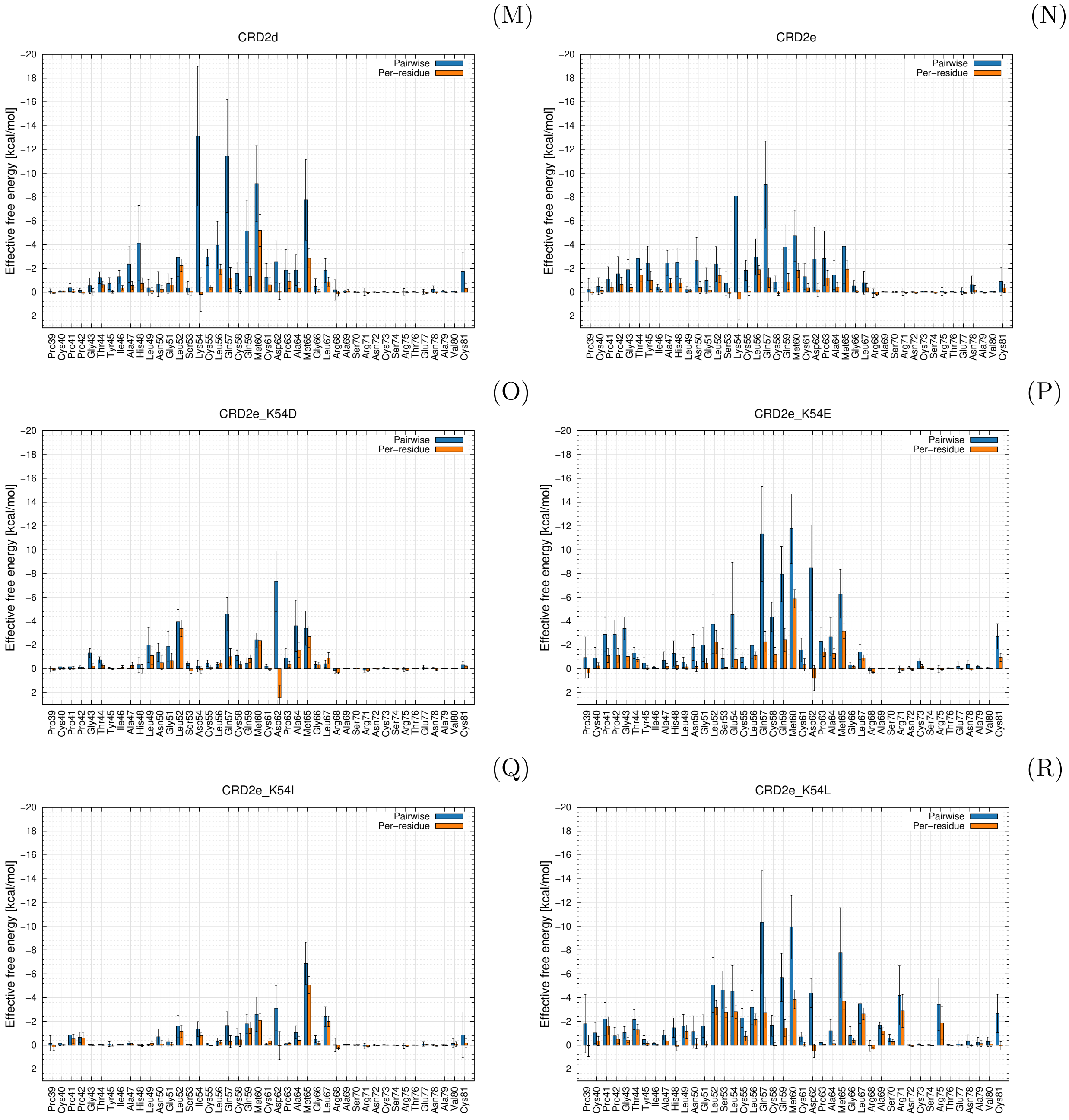

Figure S9: continued: MM-GBSA effective binding energy decomposition results for: (M) CRD2d, (N) CRD2e, (O) CRD2e_K54D, (P) CRD2e_K54E, (Q) CRD2e_K54I, (R) CRD2e_K54L variants

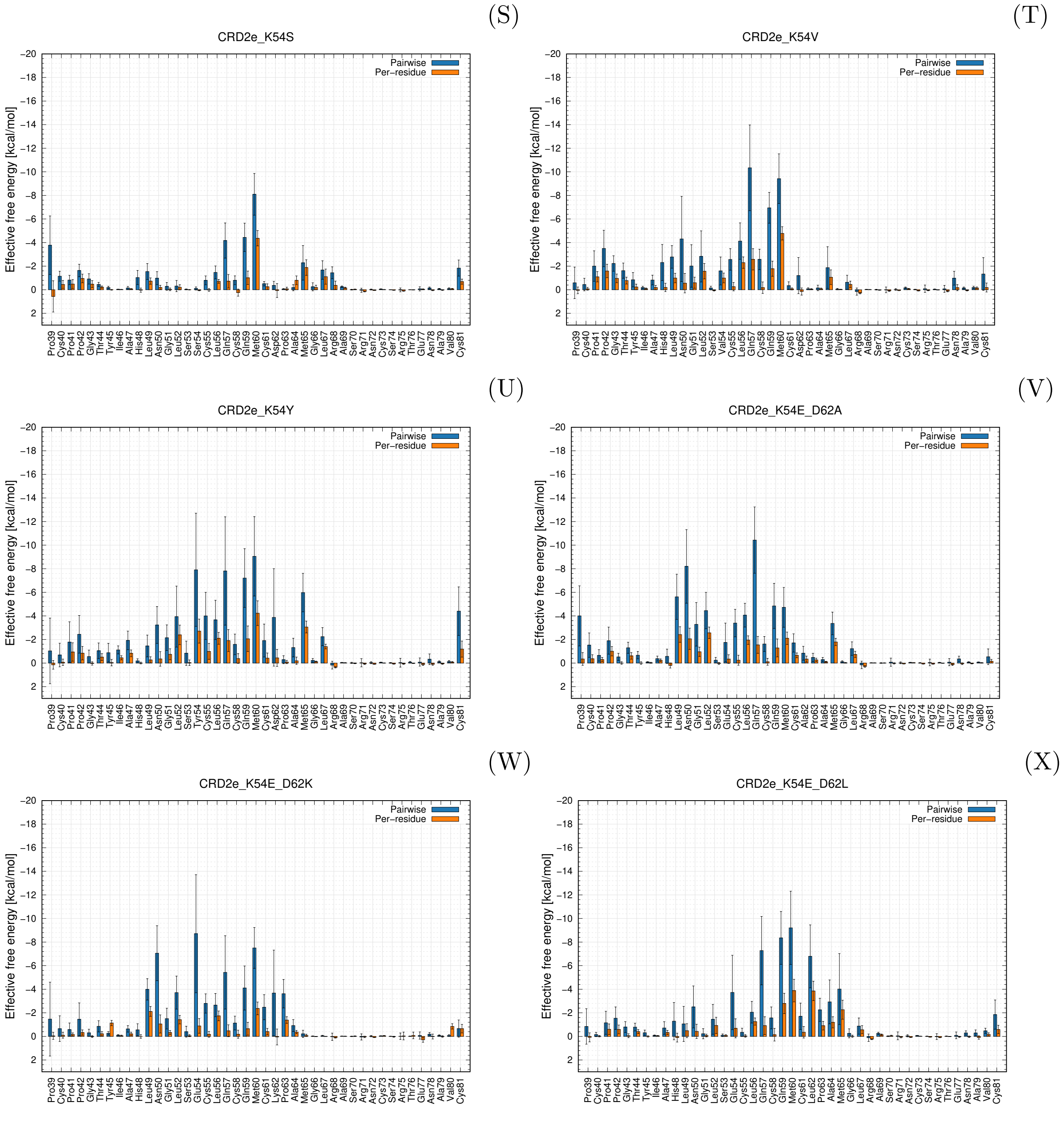

Figure S9: continued: MM-GBSA effective binding energy decomposition results for: (S) CRD2e_K54S, (T) CRD2e_K54V, (U) CRD2e_K54Y, (V) CRD2e_K54E_D62A, (W) CRD2e_K54E_D62K, (X) CRD2e_K54E_D62L variants

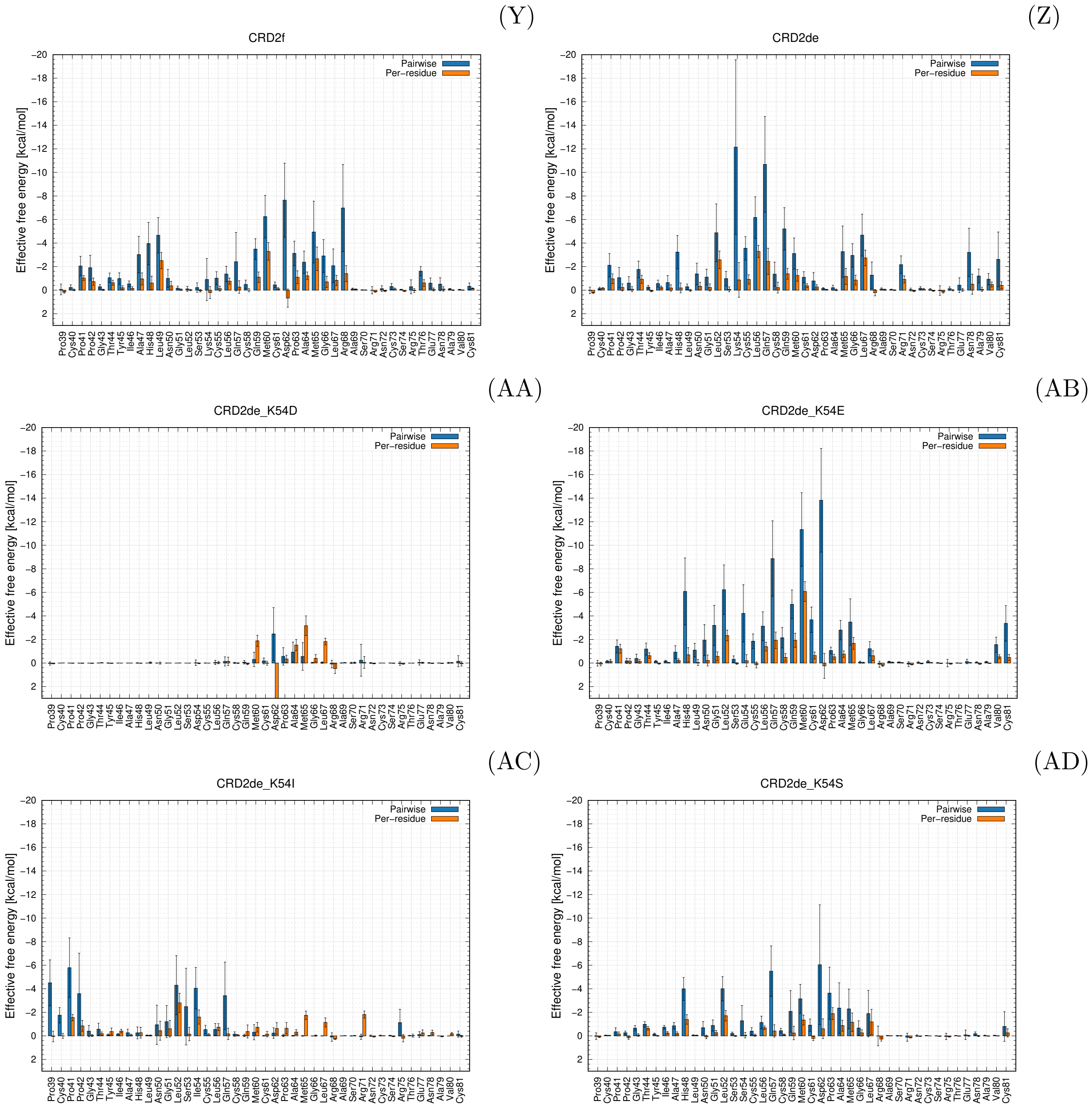

Figure S9: continued: MM-GBSA effective binding energy decomposition results for: (Y) CRD2f, (Z) CRD2de, (AA) CRD2de_K54D, (AB) CRD2de_K54E, (AC) CRD2de_K54I, (AD) CRD2de_K54S variants

(AE)

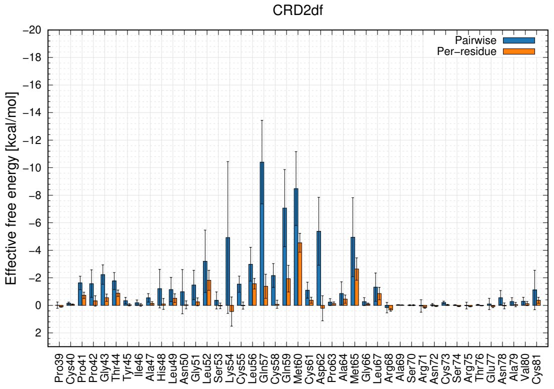

(AF)

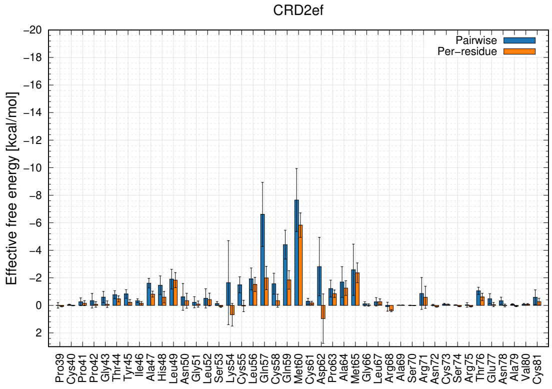

(AG)

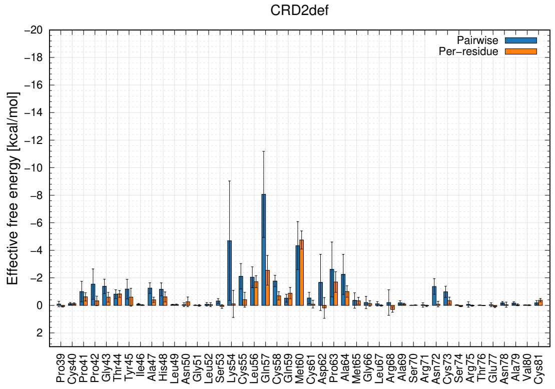

(AH)

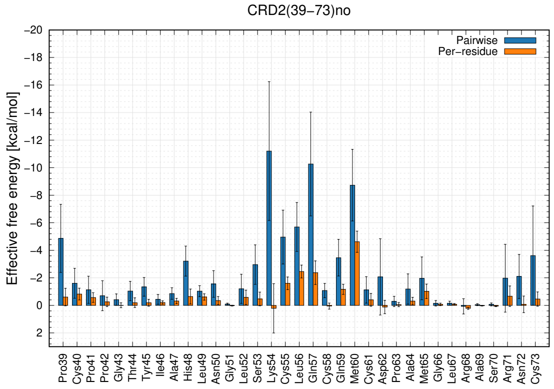

(AI)

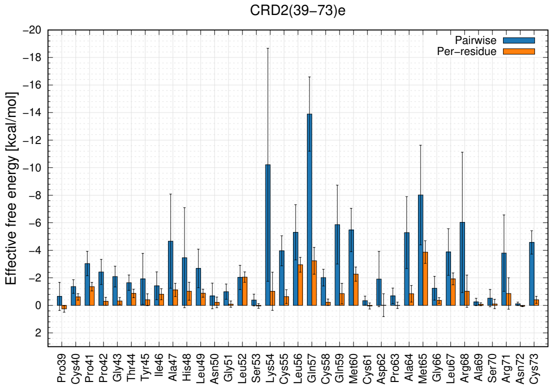

(AJ)

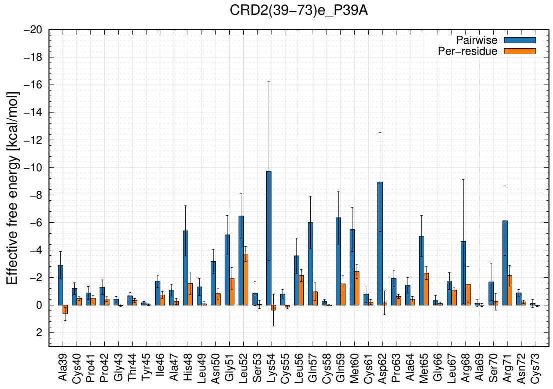

Figure S9: continued: MM-GBSA effective binding energy decomposition results for: (AE) CRD2df, (AF) CRD2ef, (AG) CRD2def, (AH) CRD2(39-73)no, (AI) CRD2(39-73)e, (AJ) CRD2(39-73)e_K54E variants

(AK)

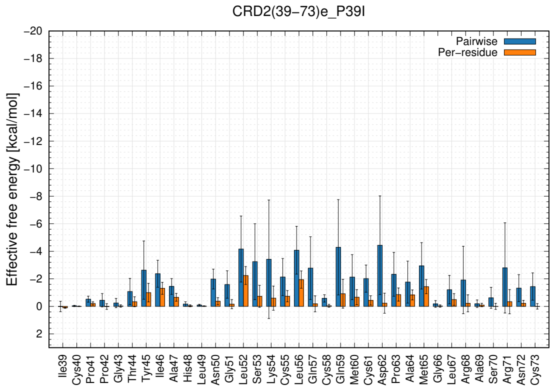

(AL)

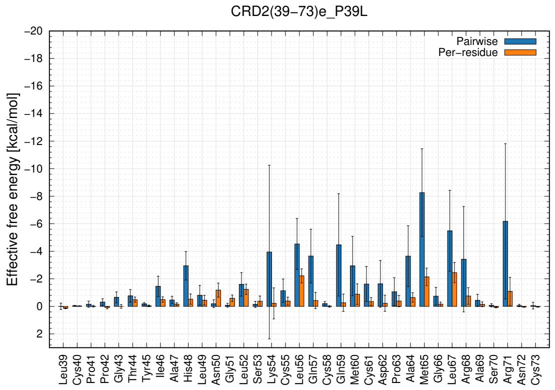

(AM)

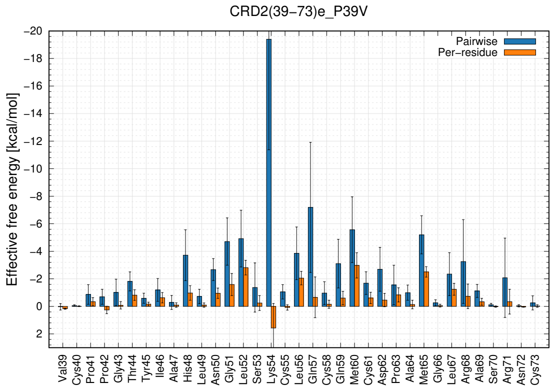

(AN)

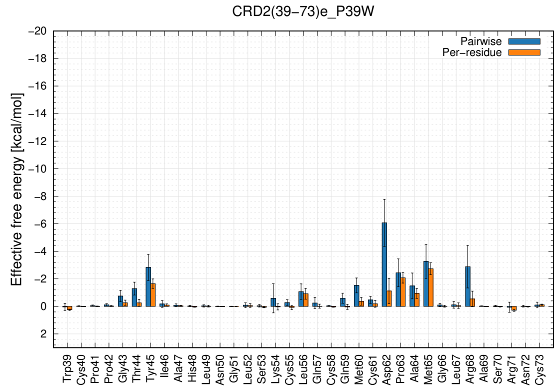

(AO)

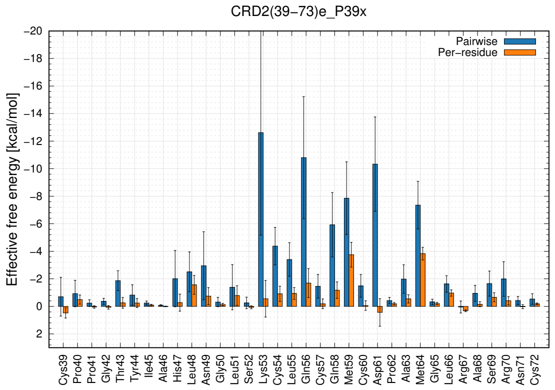

(AP)

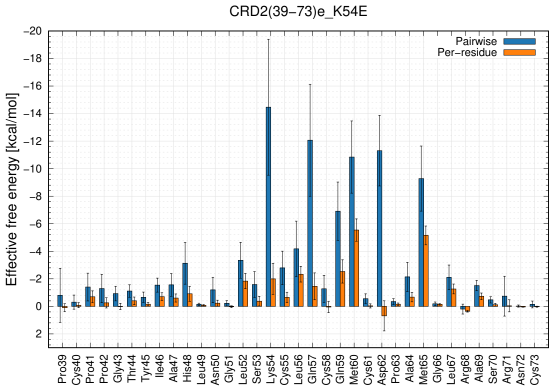

Figure S9: continued: MM-GBSA effective binding energy decomposition results for: (AK) CRD2(39-73)e_P39I, (AL) CRD2(39-73)e_P39L, (AM) CRD2(39-73)e_P39V, (AN) CRD2(39-73)e_P39W, (AO) CRD2(39-73)e_P39x, (AP) CRD2(39-73)e_K54E variants

(AQ)

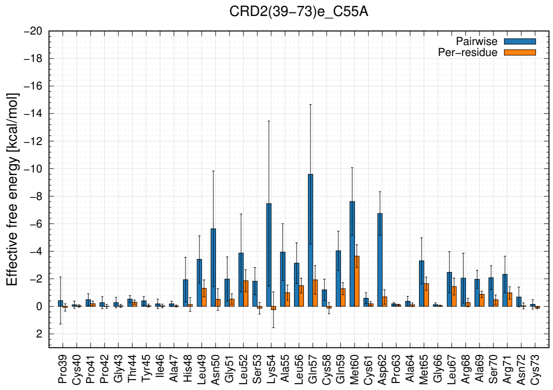

(AR)

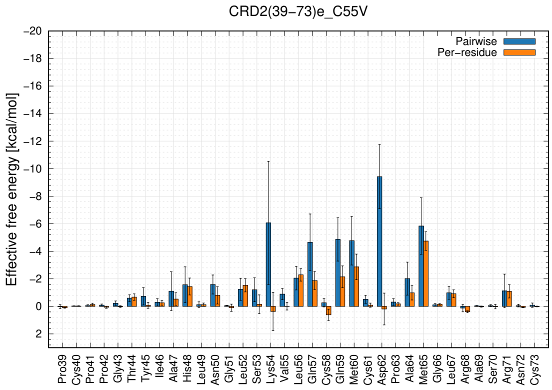

(AS)

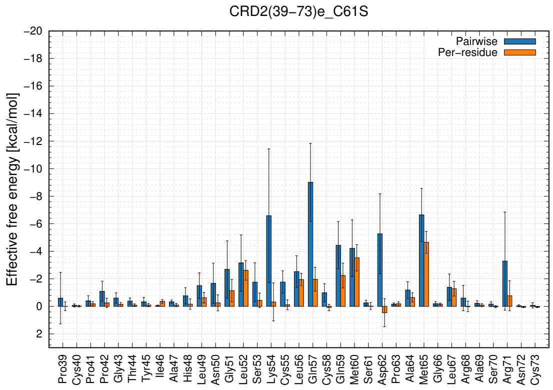

(AT)

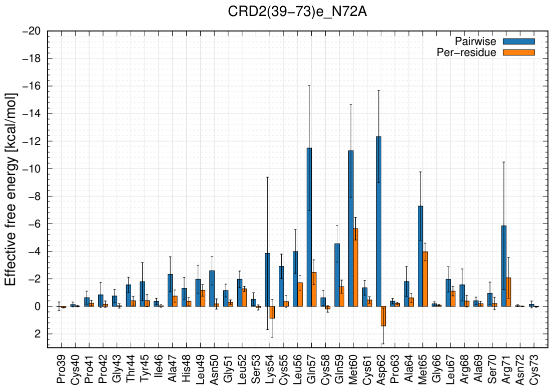

(AU)

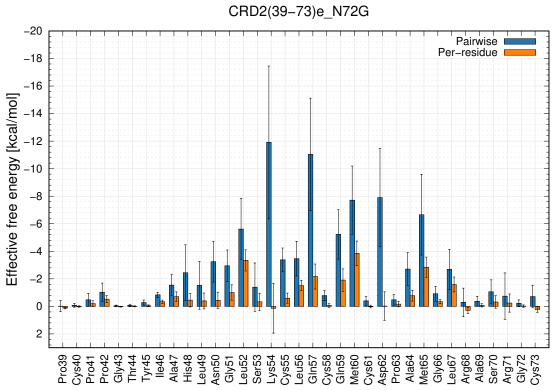

(AV)

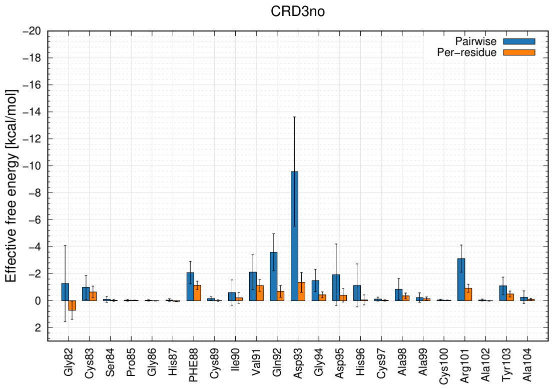

Figure S9: continued: MM-GBSA effective binding energy decomposition results for: (AQ) CRD2(39-73)e_C55A, (AR) CRD2(39-73)e_C55V, (AS) CRD2(39-73)e_C61S, (AT) CRD2(39-73)e_N72A, (AU) CRD2(39-73)e_N72G, (AV) CRD3no variants

(AW)

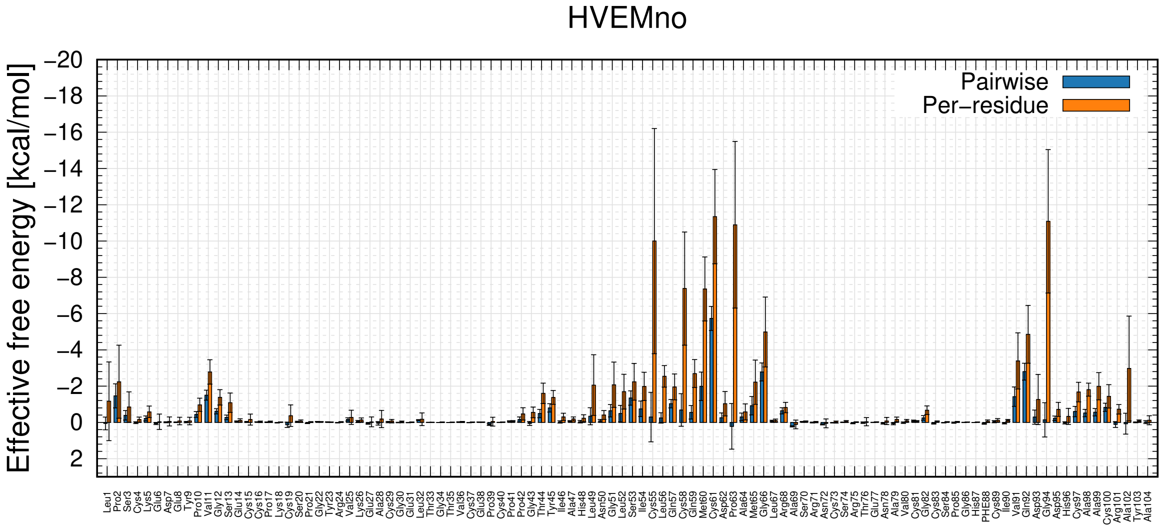

(AX)

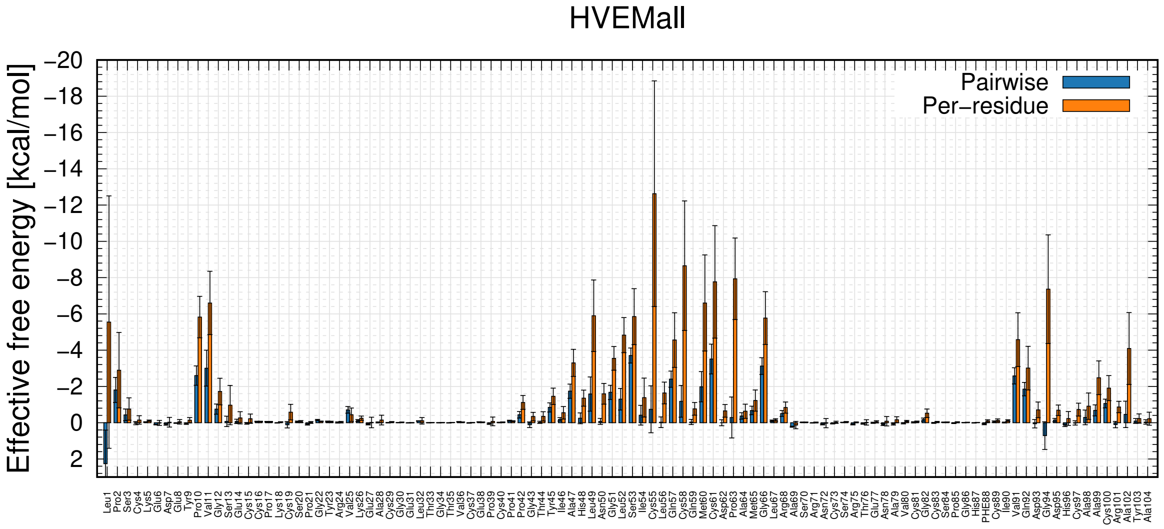

Figure S9: continued: MM-GBSA effective binding energy decomposition results for: (AW) HVEMno variant, (AX) HVEMall

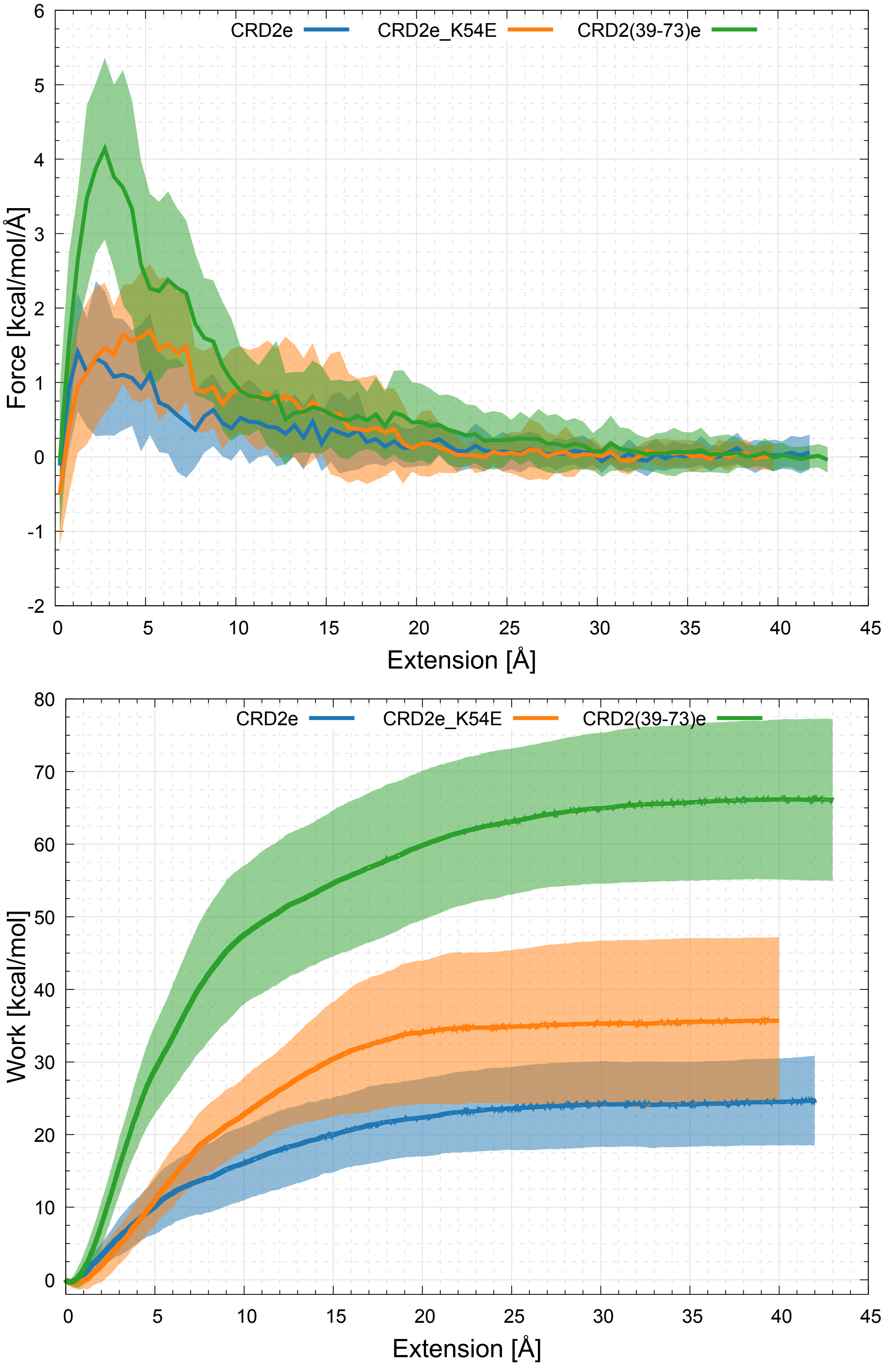

Figure S10: Plots of force and work averaged over 25 SMD trajectories (solid lines) with the corresponding standard deviation (SD) values (shown as semi-transparent areas) for HVEM variants interacting with the LIGHT trimer. Simulations were run until full dissociation was achieved.

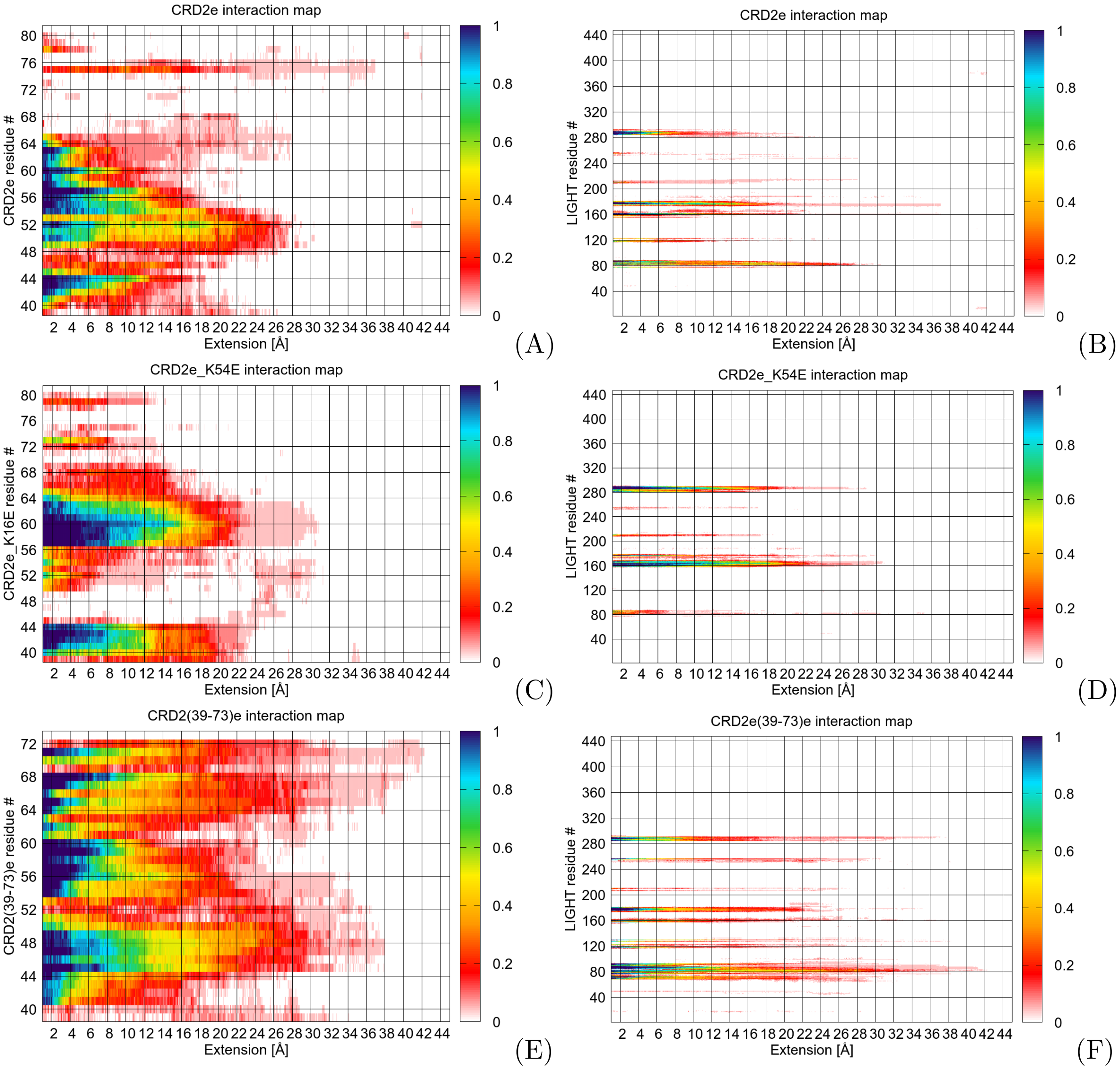

Figure S11: Contact heatmap as a function of residue index and extension (relative difference of the distance between centers of mass of HVEM variant and LIGHT trimer) averaged over 25 SMD simulations for HVEM variants with the LIGHT trimer (left panels: A, C, E), and LIGHT amino-acid residues with HVEM variants (right panels: B, D, F). The values are normalized in the range of 0-1.

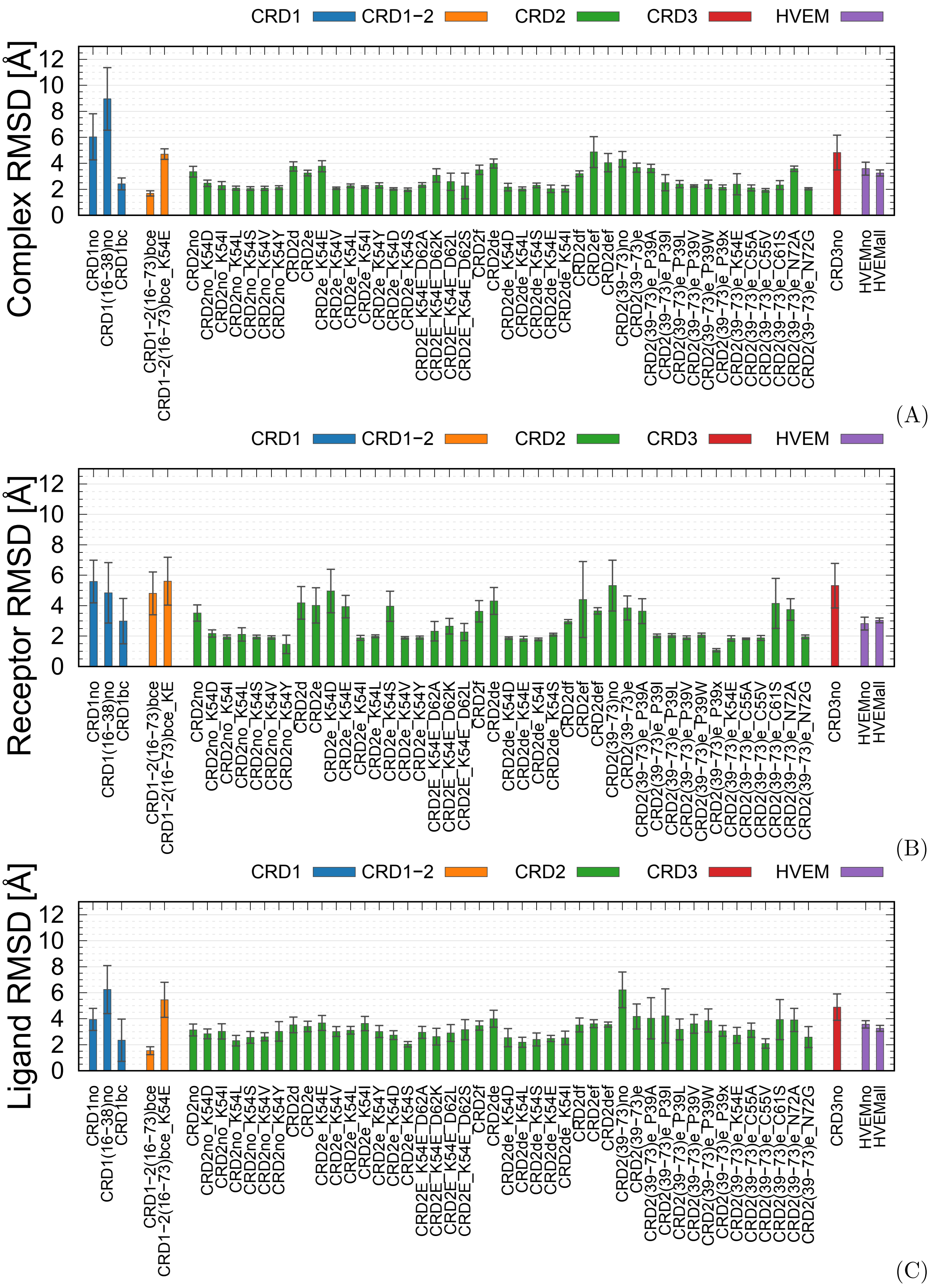

Figure S12: Bar plots of the strongly interacting complexes ($\Delta$G>50kcal/mol) of the structural properties: (A) Complex C$\alpha$RMSD, (B) Receptor C$\alpha$RMSD, and (C) Ligand C$\alpha$RMSD values averaged for 3 trajectories for the last 20 ns of simulation. For the RMSD calculation, an initial conformation adapted from the PDB file was used as a reference.

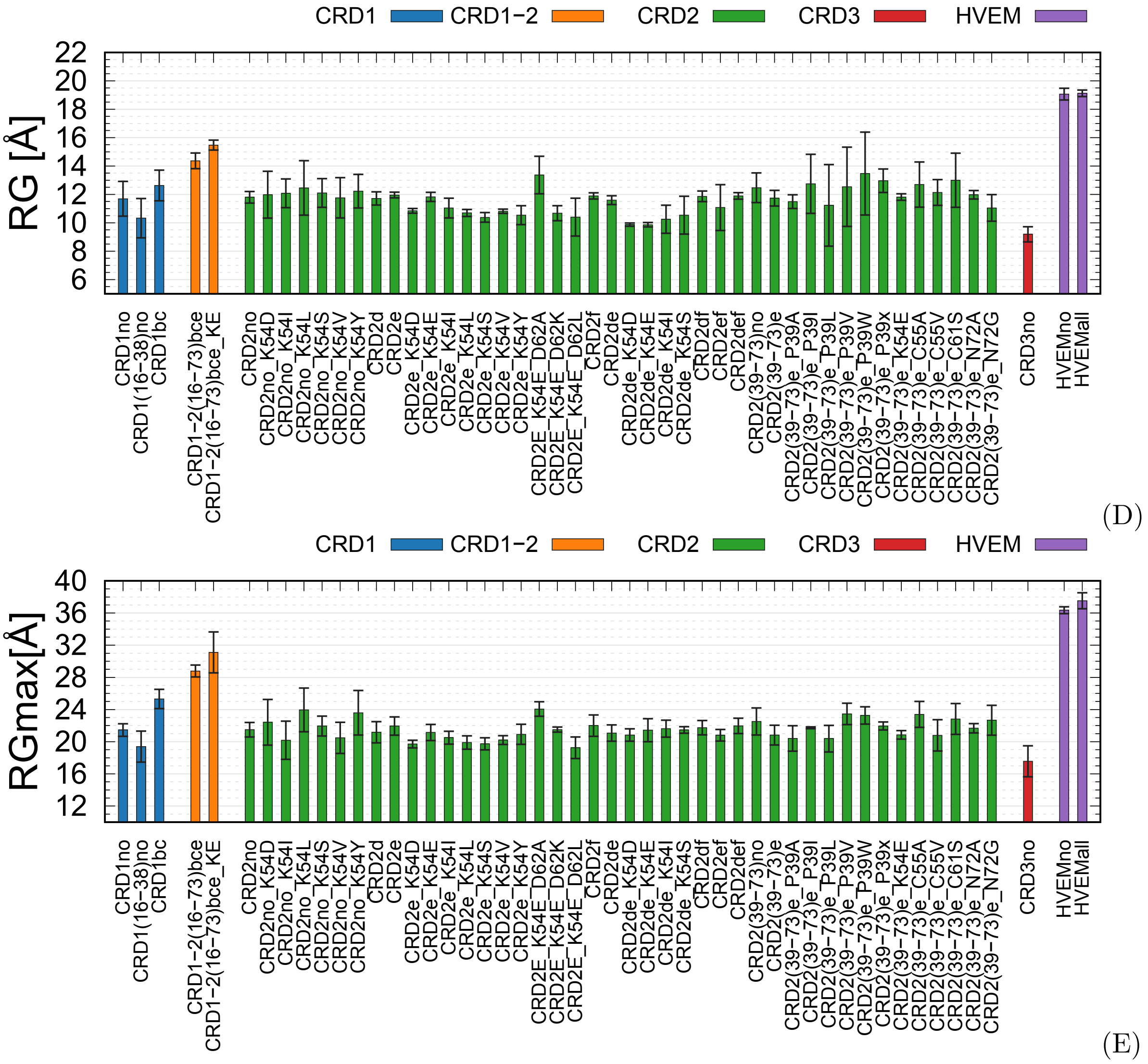

Figure S12: continued: Bar plots of the strongly interacting complexes ($\Delta$G>50kcal/mol) of the structural properties: (D) RG and (E) RGmax values averaged for 3 trajectories for the last 20 ns of simulation. For the RMSD calculation, an initial conformation adapted from the PDB file was used as a reference.

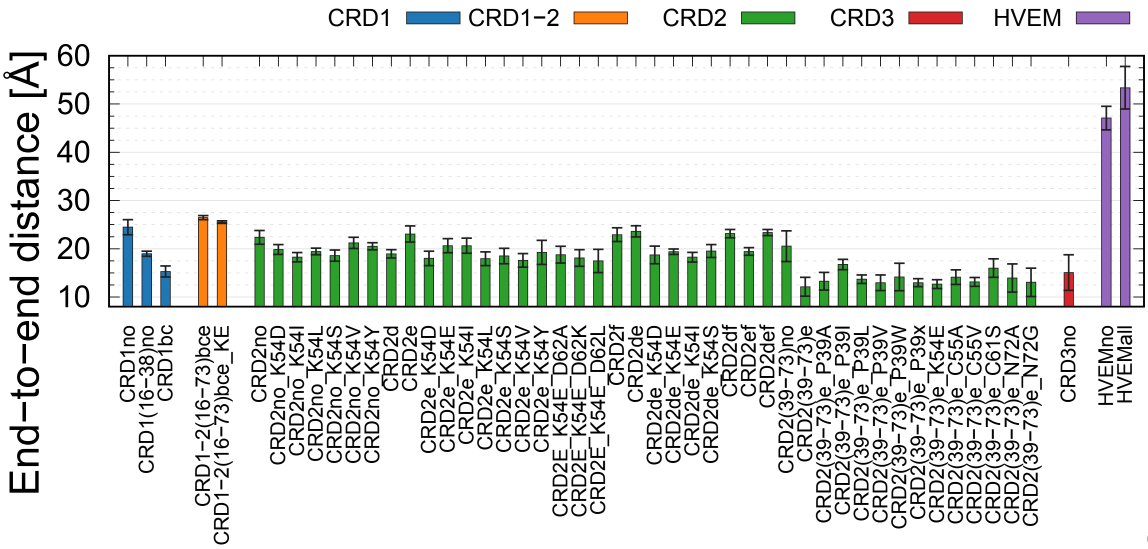

(F)

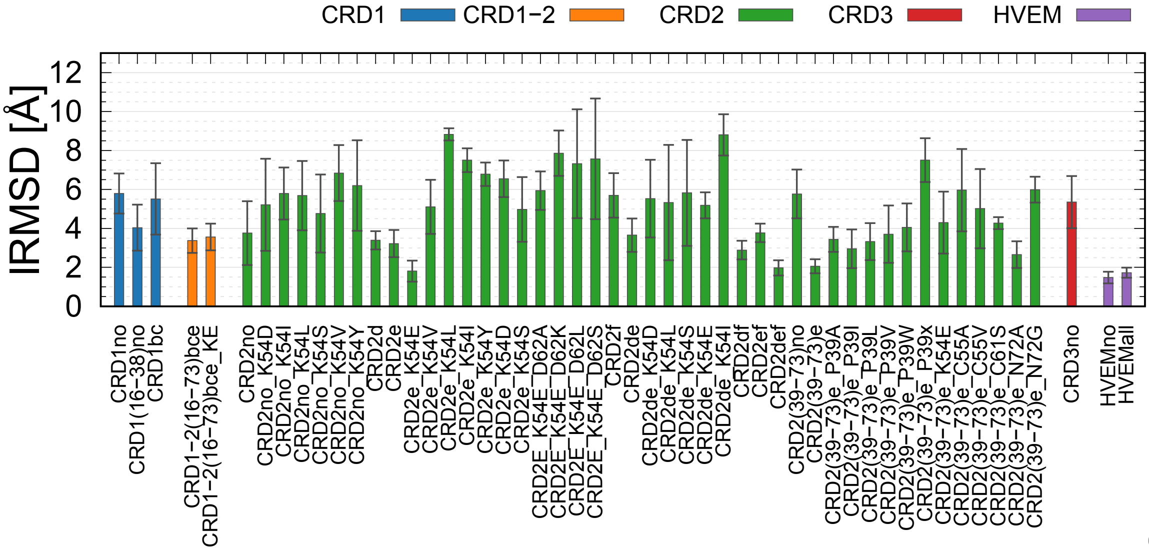

(G)

Figure S12: continued: Bar plots of the strongly interacting complexes ($\Delta G > 50$kcal/mol) of the structural properties: (F) End-to-end distance and (G) lRMSD values averaged for 3 trajectories for the last 20 ns of simulation. For the RMSD calculation, an initial conformation adapted from the PDB file was used as a reference.
